# Charting cortical-layer specific area boundaries using Gibbs’ ringing attenuated T1w/T2w-FLAIR myelin MRI

**DOI:** 10.1101/2024.09.27.615294

**Authors:** Joonas A. Autio, Akiko Uematsu, Takuro Ikeda, Takayuki Ose, Yujie Hou, Loïc Magrou, Ikko Kimura, Masahiro Ohno, Katsutoshi Murata, Tim Coalson, Henry Kennedy, Matthew F. Glasser, David C. Van Essen, Takuya Hayashi

**Affiliations:** Laboratory for Brain Connectomics Imaging, RIKEN Center for Biosystems Dynamics Research, Kobe, Japan; Université Lyon, Université Claude Bernard Lyon 1, Inserm, Stem Cell and Brain Research Institute U1208, 69500 Bron, France; Center for Neural Science, New York University, New York, NY, United States; Siemens Healthcare K.K. Tokyo, Japan; Department of Neuroscience, Washington University School of Medicine, St Louis, Missouri USA; Institute of Neuroscience, State Key Laboratory of Neuroscience, Chinese Academy of Sciences (CAS) Key Laboratory of Primate Neurobiology, Shanghai 200031, China; Department of Radiology, Washington University School of Medicine, St Louis, Missouri, USA

**Author notes:** **Corresponding author** Joonas A. Autio, Laboratory for Brain Connectomics Imaging, RIKEN Center for Biosystems Dynamics Research, 6-7-3 MI R&D Center 3F, Minatojima-minamimachi, Chuo-ku, Kobe 650-0047, Japan.

**Keywords:** Primate, Macaque, Myelin, Layer, Gibbs’ artefact

## Abstract

Cortical areas have traditionally been defined by their distinctive layer cyto- and/or myelo- architecture using postmortem histology. Recent studies have delineated many areas by measuring overall cortical myelin content and its spatial gradients using the T1w/T2w ratio MRI in living primates, including humans. While T1w/T2w studies of areal transitions might benefit from using the layer profile of this myelin-related contrast, a significant confound is Gibbs’ ringing artefact, which produces signal fluctuations resembling cortical layers. Here, we address these issues with a novel approach using cortical layer thickness-adjusted T1w/T2w-FLAIR imaging, which effectively cancels out Gibbs’ ringing artefacts while enhancing intra-cortical myelin contrast. Whole-brain MRI measures were mapped onto twelve equivolumetric layers, and layer-specific sharp myeloarchitectonic transitions were identified using spatial gradients resulting in a putative 182 area/subarea partition of the macaque cerebral cortex. The myelin maps exhibit notably high homology with those in humans, suggesting cortical myelin shares a similar developmental program across species. Comparison with histological Gallyas myelin stains explains over 80% of the variance in the laminar T1w/T2w-FLAIR profiles, substantiating the validity of the method. Altogether, our approach provides a novel, noninvasive means for precision mapping layer myeloarchitecture in the primate cerebral cortex, advancing the pioneering work of classical neuroanatomists.

## Introduction

For over a century, myelin staining has been a fundamental technique for delineating cortical layers and parcellating areas of the cerebral cortex (Baillarger, 1840; Vogt and Vogt, 1919; Vogt, 1910; Vogt 1912; Hopf 1968, 1969, Nieuwenhuys et al. 2013, 2017). Myelin enwraps fundamental circuit motifs such as cortico-thalamo-cortical projections, the bands of Baillarger, and inter-areal feedforward and feedback connections, which are critical for cortical functions (Jones and Burton 1976, Felleman and Van Essen 1991). Although these circuits are fundamentally linked with information flow in the cerebral cortex (Markov et al., 2013), the link between variations in cortical layer myelination and areal position within the cortical hierarchy remains poorly understood (Felleman and Van Essen 1991, Burt et al., 2018, Harris et al., 2019).

There is a growing interest in differentiating cortical areas using non-invasive magnetic resonance imaging (MRI). Initially, these efforts primarily focused on regional differences in overall myelin content. One way to improve the sensitivity to the intra-cortical myelin is the mapping based on the ratio between T1-weighted (T1w) and T2-weighted (T2w) images (Glasser and Van Essen 2011; Glasser et al., 2014; 2016), aimed at improving the contrast-to-noise ratio (CNR) for intracortical myelin while mitigating undesirable contrast mechanisms (e.g., B_1_ receive bias-field). This approach enabled noninvasive mapping of cortical content and delineation of cortical areas in humans (Glasser and Van Essen 2011; Glasser et al., 2016) and non-human primates (Glasser et al., 2014, Donahue et al., 2018, Ikeda et al., 2022). However, a gap exists between overall MR myelin measures and the classical determination of laminar patterns, and not much is known about the cortical layer distribution of the T1w/T2w ratio (Eickhoff et al., 2005, Petracca et al., 2020, Li et al., 2019, Wei et al., 2022). Therefore, noninvasive multi-modal parcellations may further benefit from collecting more fine-grained information on cortical layer myeloarchitecture.

Since seminal postmortem laminar MRI studies (Clark et al., 1992, Fatterpekar et al., 2002), an increasing number of high-resolution MRI studies have pursued whole-brain cortical layer mapping in vivo (Dinse et al., 2015; Large et al., 2016; McColgan et al., 2021; Petracca et al., 2020; Waehnert et al., 2014, Fracasso et al., 2016, Tomer et al., 2022, Avram et al., 2021). These studies have revealed layer-dependent contrasts in T1w (Stuber et al., 2014), T1-mapping (Lifshits et al., 2018, Tomer et al., 2022), T2w susceptibility-weighted (Fukunaga et al., 2010) and diffusion-weighted (DW) (Assaf, 2019) imaging. Yet a hemisphere-wide cortical surface mapping of laminar transitions has not yet been fully realised due to prominent in vivo MRI challenges, including involuntary motion, resolution limitations relative to layer thicknesses (Lifshits et al., 2018), errors in bias-field correction (Glasser et al., 2011), and inter-subject variability which is more prominent in humans than in non-human primates (NHPs) (Coalson et al., 2018, Hayashi et al., 2021). Consequently, demonstration of layer differences between cortical areas has primarily been limited to relatively small volumes of cortical ex vivo samples.

Another significant challenge in MRI of the highly convoluted cerebral cortex is to distinguish between ‘true’ laminar MR contrasts (e.g. myelin, iron, and blood) and imaging artefacts oriented parallel to the cortical sheet. In particular, a well-known Gibbs’ ringing artefact (also known as truncation-artefact) occurs in signal intensity transitions that are spatially sharp and large in magnitude, such as that between cerebro-spinal fluid (CSF) and cortical grey matter (CGM). These may manifest as spurious signal-intensity damped oscillations parallel to the cortical surface, due to the Fourier transformation during image reconstruction (Gibbs, 1899). All MR image types having sharp image contrast transitions are affected by Fourier image reconstruction artefacts (Dinse et al., 2015; Perrone et al., 2015; Veraart et al., 2016). Furthermore, these artefacts can be affected by cortical folding, owing to constructive or destructive interference across opposed banks of a cortical sulcus. One way to reduce ringing artefacts is to increase spatial resolution, but this is limited by the need for a high signal-to-noise ratio (SNR), and artefacts can nonetheless persist proximal to the sharp signal-intensity transition zones. Several post-processing methods have been proposed to overcome the ringing artefacts, including image filters (e.g. hamming window, Lanczos σ-approximation) (Jerri, 2000), total variation constrained data extrapolation (Block et al., 2008; Veraart et al., 2016), and local sub-voxel shifts in conjunction with saddle-point *k*-space filters (Kellner et al., 2016). However, image filters predictably blur images and reduce effective image resolution, while methods based on signal-intensity variation clean-up are not designed to selectively attenuate ringing artefacts and they may also reduce genuine layer-specific MR signal variations.

Here, we present a new laminar myelin mapping strategy in macaque monkeys in vivo that provides two key advances relative to the conventional T1w/T2w approach (Glasser et al., 2011, 2013, 2016). One involves higher spatial resolution than the 0.5 mm (0.125 mm^3^) voxels commonly used for macaque structural scans (e.g., Donahue et al. 2016; Autio et al., 2021). We scaled image acquisition according to median cortical thickness (1.9 mm in macaque) divided by the standard number of layers in the isocortex (1.9 mm/6 layers = 0.32 mm; 0.033 mm^3^) so that each layer has at least one independent voxel. The second was to replace the conventional T2w-SPACE scan, in which white matter (WM) is dark, CGM is intermediate in intensity, and CSF intensity is very high, with a T2w-FLAIR scan, which has similar intensity contrast as T2w-SPACE scan but low CSF intensity. This makes the CSF-CGM dark-to-light contrast similar to that in a conventional T1w MPRAGE scan but keeps the contrast between CGM and WM opposite in polarity between T2w-FLAIR and T1w. We demonstrate that the prominent Gibbs’ ringing artefact at the CSF/CGM border largely cancels out when taking the T1w and T2w-FLAIR ratio. We quantitatively validate this methodology using a postmortem myeloarchitectonic ground-truth derived from modified Gallyas stains and demonstrate cortical layer-specific myeloarchitectonic area boundaries with unprecedented accuracy. Finally, we explore the link between these laminar myeloarchitectonic features and hierarchical organisation of the cerebral cortex.

## Results

### Overcoming Cortical Layer-Like Gibbs’ Ringing Artefacts Using T1w/T2w-FLAIR for Anatomically Informative Layer MRI

Representative zoomed views of median cortical layer thickness adjusted (320 µm isotropic) T1w, T2w, and T2w-FLAIR images are shown in Figure 1A. The images exhibit pronounced contrast differences along the cortical radial axis. However, the precise nature of these layer-like structures, particularly evident in the T2w image, remains ambiguous as to whether they represent genuine anatomical layers or Gibbs’ ringing artefacts arising from the Fourier transformation in the MR image reconstruction. Indeed, simulation of Gibbs’ ringing artefact predicts the strongest signal oscillations on the superficial cortical layers at a spatial frequency of cycle / (2 × voxels) (= 640 µm) (Fig. 1B, C). The spatial frequency of this artefact was experimentally verified by acquiring T2w images using different resolutions (Supp. Fig. 1A, B). Importantly, the phase of Gibbs’ ringing artefact differs between the two imaging modalities: T1w and T2w images exhibit opposing-phase ringing artefacts (Fig. 1C; blue and red lines, respectively). Consequently, T1w and T2w image ratio amplifies the Gibbs’ ringing artefact, posing a substantial challenge in distinguishing between artefacts and genuine anatomical layers (Fig. 1C; purple line).

**Figure 1.**
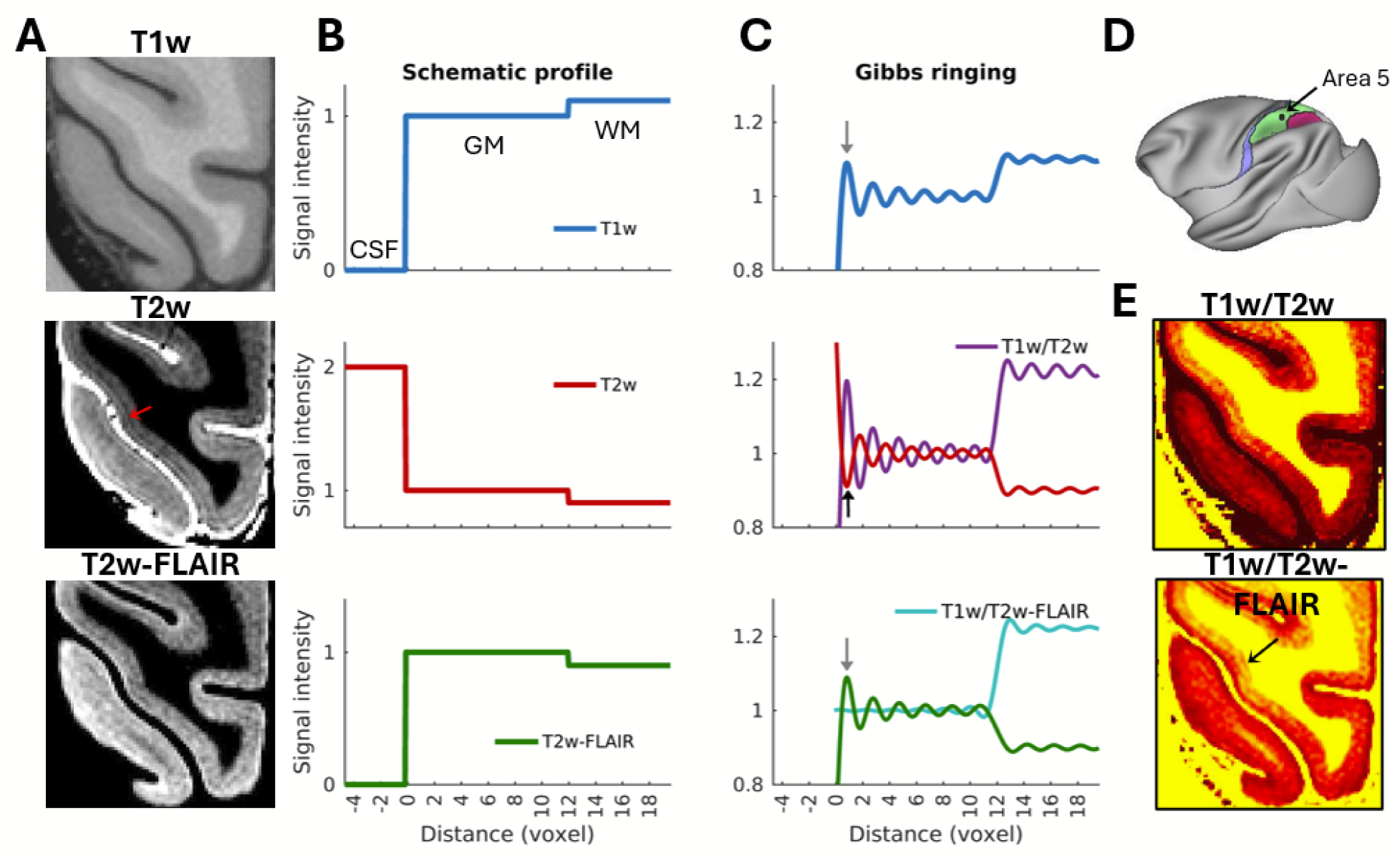
Attenuating Gibbs’ Ringing Artefact using T1w and T2w-FLAIR ratio. **(A)** Representative magnified view of cortex displaying T1w, T2w and T2w-FLAIR contrasts. Images are a combination of true anatomical contrast and Gibbs ringing artefacts. In the T2w image, the red arrow denotes a distinct Gibbs’ ringing artefact which resembles cortical layers. **(B, C)** Simulation of Gibbs’ ringing artefact. Sharp positive signal intensity transitions cause an initial overshoot followed by sinc-pattern decay (grey arrows), while negative signal-intensity transitions cause an undershoot (black arrow) in the outermost voxel in grey matter (GM). Note that the phase of Gibbs’ ringing artefact is reversed by fluid-attenuated inversion-recovery (FLAIR; bottom panel, grey arrow). The magnitude of truncation artefact varies based on signal intensity difference between tissues (e.g. contrast between CSF, GM and WM). **(C)** The in-phase ringing artefacts can be effectively attenuated by dividing T1w by T2w-FLAIR (cyan line; bottom panel). In contrast, T1w divided by T2w amplifies the ringing artefact due to the opposing phase of the Gibbs’ overshoot (purple line; middle panel). **(D)** Location of magnified view. Area 5, known for the relatively wide inner and narrow external band of Baillarger, is highlighted in green. **(E)** T1w/T2w (upper panel) and T1w/T2w-FLAIR (bottom panel) exhibit notable image contrast differences across cortical layers.

Recognizing that the largest T1w/T2w Gibbs’ ringing amplification originates from signal-intensity differences between cranium, CSF and cortex, a simple and efficient method to reduce the amplitude and number of ringing artefacts is to use T2w with fluid-attenuated inversion recovery (T2w-FLAIR) (Supp. Fig. 1C). Subsequently, the T1w and T2w-FLAIR images produce *in-phase* Gibbs’ ringing artefact at the CSF and CGM interface (Fig. 1B), and therefore T1w/T2w-FLAIR ratio effectively attenuates the remaining ringing artefacts (Fig. 1C, Supp. Fig. 1F), irrespective of their precise locations, subvoxel shifts and superimposed ringing artefacts (Kellner et al., 2016). Finally, signal-intensity transition between CGM and WM is much smaller in amplitude and more gradual than at the CGM and CSF interface due to the penetrating myelinated fibres from white matter. Thus, Fourier expansion (by mathematical definition) more accurately captures the WM-CGM transition yielding negligible small Gibbs’ ringing artefact. Overall, these features represent a novel experimental approach to effectively attenuate the Gibbs’ ringing artefact for improved cortical layer myeloarchitectonic imaging (for more detailed description, see *SI Data Acquisition Strategy - Attenuating of Gibbs’ Ringing Artefact*).

In each subject, the Gibbs’ ringing artefact attenuated T1w/T2w-FLAIR volume image, was mapped to twelve laminar surfaces estimated using an equivolume model (for representative surfaces, see Supp. Fig. 2) (Bok 1929, Van Essen 1980; Waehnert et al. 2014; Wagstyl et al., 2018) and corrected for cortical curvature and B_0_ orientation bias. Throughout the text, we refer to the equivolumetric layers (EVLs) using Arabic numbers and lower-case d/s (deep/superficial) ranging from white matter, EVL6d, 6s, 5d, 5s, …, 1s to pial surface). In contrast, we refer to anatomical cortical layers using layer and Arabic number (e.g. L4b), or roman numerals (e.g. layer IVb). This nomenclature is explicitly intended to distinguish the EVLs from genuine anatomical layers which vary in thickness and may also differentially vary on gyral and sulcal folds, but see (Wagstyl et al., 2018).

The EVL T1w/T2w-FLAIR profiles revealed non-monotonic and fine-grained myeloarchitectonic details throughout the cortical mantle. In general, the T1w/T2w-FLAIR ratio was highest adjacent to white matter and reduced towards more superficial cortical layers. Laminar T1w/T2w-FLAIR profiles exhibited signal intensity peaks (e.g. bands of Baillarger) that were systematically observed across most of the neocortex (Supp. Fig. 3C). For example, the densely myelinated external band of Baillarger (e.g. stria of Gennari) in V1 was clearly distinguishable at laminar T1w/T2w-FLAIR profiles (Fig. 2A arrow, D). These peak locations are in contrast to Gibbs’ ringing artefacts, which are most pronounced adjacent to the pial surface (Fig. 1C, Supp. Fig. 1A, D).

**Figure 2.**
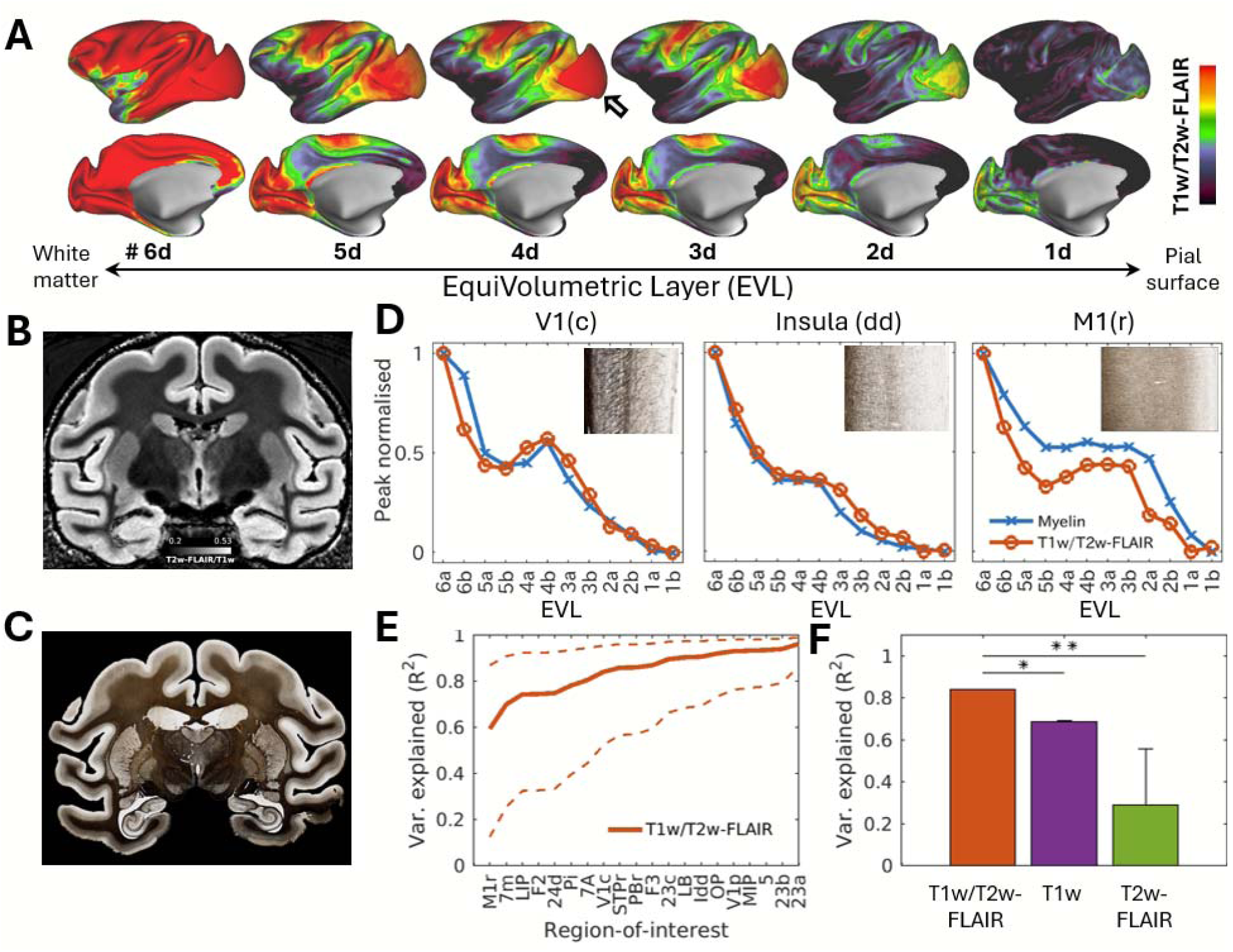
Validating Laminar Myelin Mapping. **(A)** Average T1w/T2w-FLAIR representative equivolumetric layers (EVLs) show that deep layers (left) exhibit relatively higher myelination in comparison to superficial layers (right) throughout the macaque cerebral cortex (N=7). The grey arrow indicates the outer band of Baillarger in V1. Representative sections from **(B)** T1w/T2w-FLAIR ratio and **(C)** Gallyas myelin histology. For visualisation purposes, the T1w/T2w-FLAIR ratio is inverted (e.g., T2w-FLAIR/T1w). **(D)** Exemplar peak normalised layer profiles of Gallyas myelin stain (blue line) and T1w/T2w-FLAIR ratio (orange line). Snippets show myelin histology from respective cortical areas. **(E)** T1w/T2w-FLAIR is highly correlated with myeloarchitecture (n=20). **(F)** Compared to T1w and T2w-FLAIR, T1w/T2w-FLAIR provided significantly improved correlations with layer myeloarchitecture (*p < 10^-4^, **p < 10^-15^, paired t-test).

We next sought to validate the equivolumetric T1w/T2w-FLAIR profiles with respect to a ‘gold-standard’ modified Gallyas myelin stain (Gallyas et al., 1979). We determined 30 regions-of-interest (ROIs) in well-stained histology and selected corresponding profiles in atlas MRI space (Fig. 2B, C). Across the ROIs, the Gallyas myelin stain and T1w/T2w-FLAIR laminar profiles were strikingly similar (Fig. 2D, E). Average correlation across ROIs between myelin histology was significantly higher in the T1w/T2w-FLAIR (R^2^ = 0.84 ± 0.09; mean ± std) in comparison to T1w (R^2^ = 0.69 ± 0.14, p < 0.0001; paired t-test, Bonferroni corrected) and T2w-FLAIR (R^2^ = 0.29 ± 0.27, p <10^-15^) indicating that T1w/T2w-FLAIR, indeed, provides improved specificity as well as sensitivity to intra-cortical myelination (Fig. 2F).

### Charting Cortical Area Boundaries using Sharp Layer-Specific Myeloarchitectonic Transitions

Having established a close correlation between T1w/T2w-FLAIR and myelin histology, we next focused on delineating cortical area boundaries using layer-specific myeloarchitectonic transitions. For this objective, we calculated gradient-ridges on each EVL (Fig. 3A, B) and explored sharp and layer-specific myeloarchitectonic transitions. For instance, in the sensorimotor cortex, the EVL3d exhibited strong myelin gradient-ridges across areas whereas EVL5d exhibited weak gradients (Fig. 3A-D). This result is in excellent agreement with Gallyas myelin stain (Fig. 3E): sensorimotor areas exhibit more pronounced differences in superficial layers and only subtle differences in the deep cortical layers. Our noninvasive determination of area boundaries were in good agreement with previous atlases (Supp. Fig. 4), albeit the area boundaries differed in the medial cortex aligning more with the view of subdivisions of the M1 (see Supplementary Discussion; Rathelot et al., 2009). Taken together, these results demonstrate that mapping layer-specific transitions may have significant advantages for delineating cortical area boundaries.

**Figure 3.**
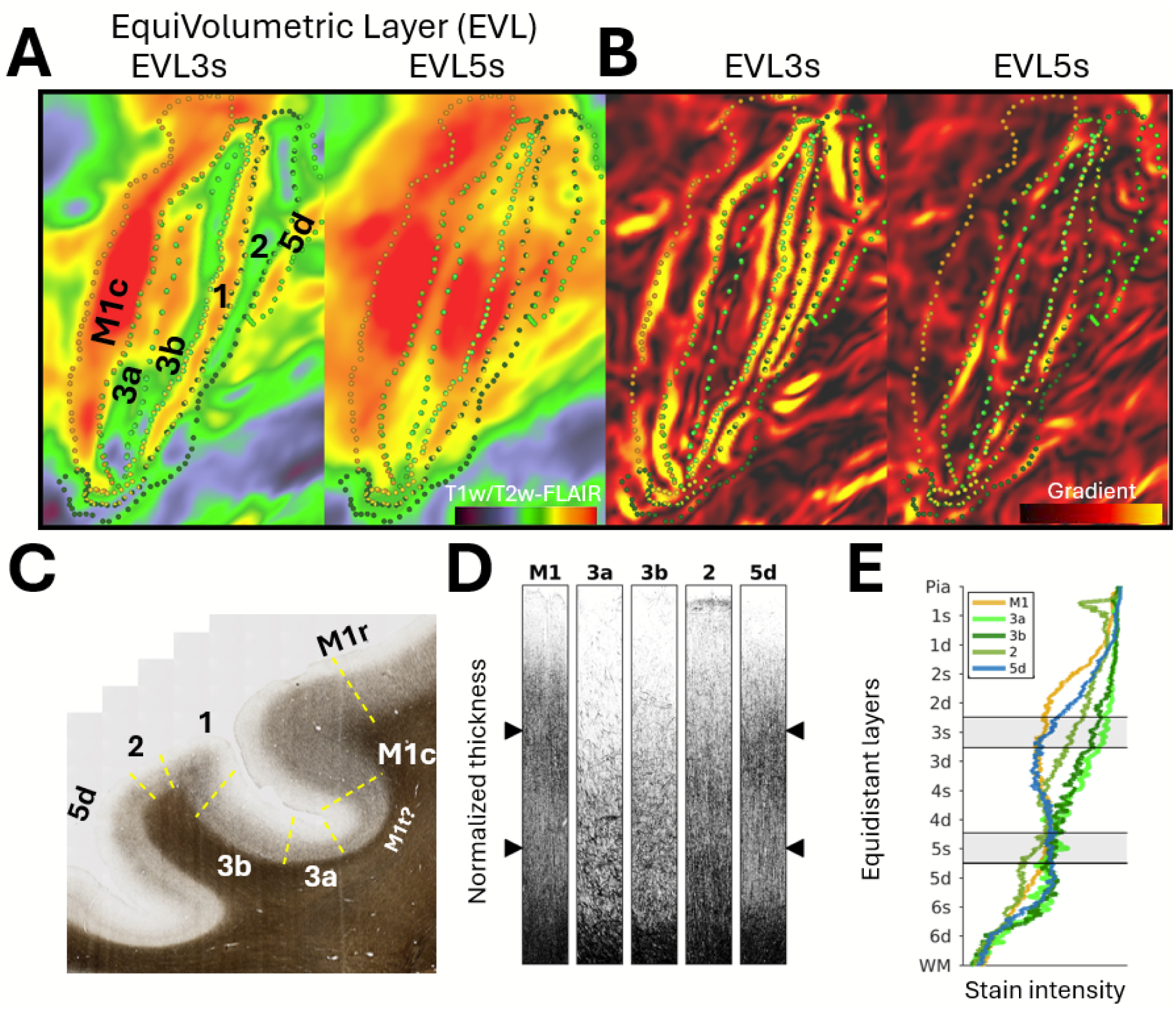
Charting cortical area boundaries using cortical layer-specific myeloarchitectonic transitions. **(A)** Zoomed view of sensorimotor cortex in representative equivolumetric layers (EVL) 3d and 5d displayed on a flat-map. **(B)** T1w/T2w-FLAIR gradients are prominent in EVL3d but less so in EVL5d. **(C)** Gallya’s myelin stain from somatomotor cortices. Yellow dashed lines indicate area boundaries. **(D)** Cortical thickness normalised laminar profiles of representative myelin stains. Notably, large myelin density differences are observed in superficial layers (e.g., EVL3s), whereas deeper layers exhibit more modest differences (e.g. EVL5s). Abbreviations: M1c: caudal Primary Motor Cortex caudal (Rathelot et al., 2009); M1t: transitory primary motor cortex (current study).

We applied this approach to delineate 182-area/subarea boundaries in the left hemisphere (Fig. 4A) and a comparable number in the right hemisphere (see BALSA). Sharp myelin gradient ridges were observed in unimodal regions such as the auditory cortex (Supp. Fig. 5), and primary interoceptive areas (Supp. Fig. 9), as well as in transmodal regions such as the middle temporal cortex (Supp. Fig. 6) and the intraparietal cortex (Supp. Fig. 7). In transmodal areas, myeloarchitectonic transitions were more pronounced in middle cortical layers (e.g. EVLs 5 and 6). The myelin gradient-ridges became progressively weaker in middle layers, making the parcellation more reliant on deep layers as the hierarchical distance from primary sensory cortices increased (Felleman and Van Essen 1991, Markov et al., 2014). For instance, orbitofrontal (Supp. Fig. 8), agranular insula (Supp. Fig. 9), and medial prefrontal cortices, which are involved in supramodal processing, were the most challenging regions to parcellate due to very weak myelin contrast, resulting in the largest uncertainties in these regions.

**Figure 4.**
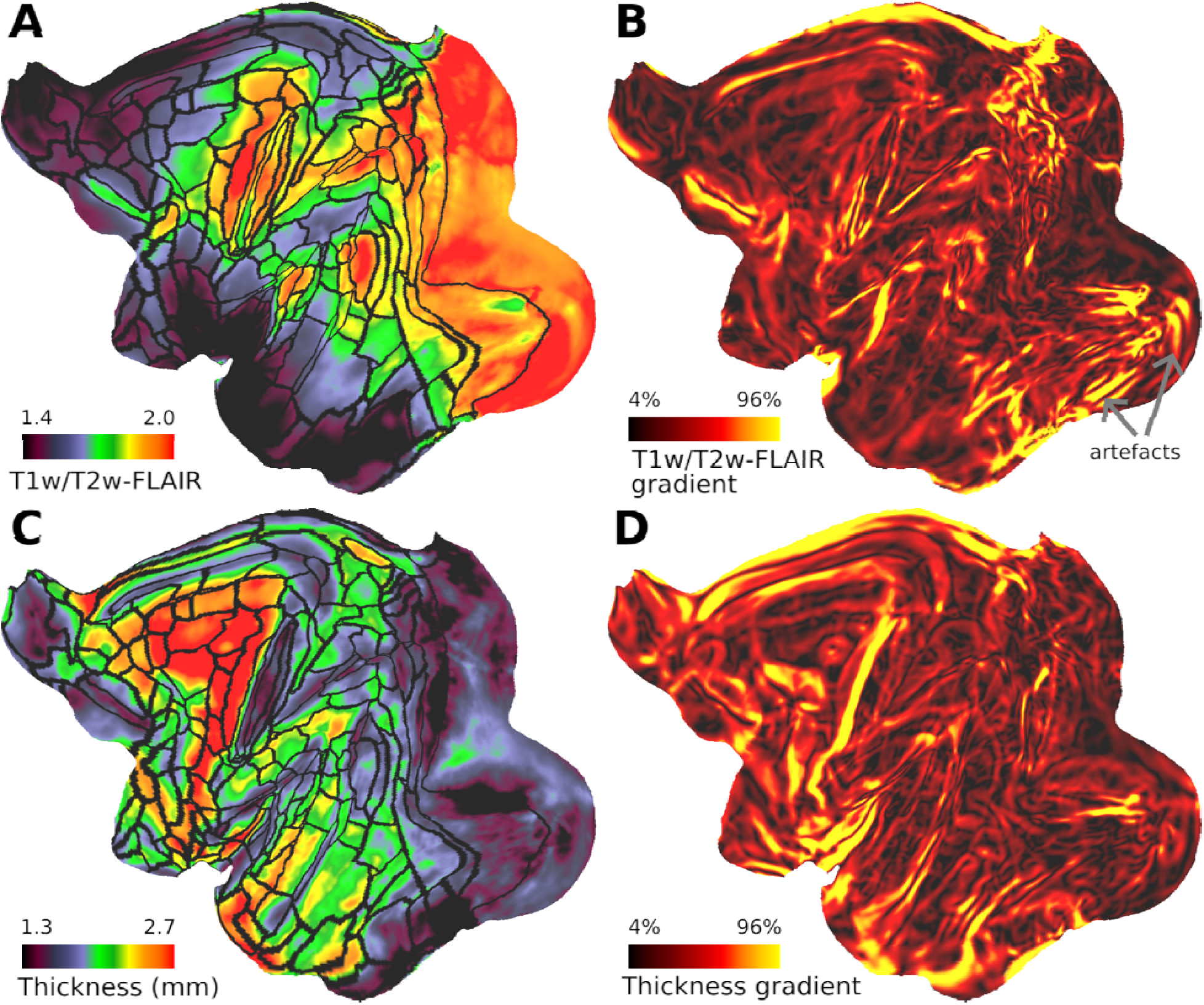
Equivolumetric Layer-specific Myelin Transitions provide Improved Sensitivity to Cortical Area Boundaries Compared to Midthickness-weighted Gradients. **(A)** Midthickness-weighted T1w/T2w-FLAIR overlaid with parcel boundaries determined using layer-specific T1w/T2w-FLAIR gradients (black contours). **(B)** Midthickness-weighted T1w/T2w-FLAIR gradients exhibit notably lower number of areal gradient-ridges due to reduced specificity to layer-specific myeloarchitectonic transitions. Grey arrows indicate artefacts. **(C)** Cortical thickness and its **(D)** gradients.

The advantage of charting cortical layer-specific myeloarchitecture over conventional midthickness-weighted myelin imaging is demonstrated in Figure 4. In comparison to EVL gradients (Fig. 4A, summarised by black lines), the midthickness-weighted gradients exhibited substantially fewer area boundaries (Fig. 4B). On the other hand, we also found that some of the borders were more robust in the midthickness–weighted myelin maps. For instance, in orbitofrontal cortices it was challenging to delineate cortical area boundaries using EVL myelin gradient maps, however, in the midthickness-weighted myelin maps the gradients appeared more robust suggesting that areal myelin differences were more evenly distributed across cortical layers (Supp. Fig. 8). Together, these results highlight the advantage of mapping cortical-layer specific myeloarchitectonic transitions as well as averaging across the layers for delineating areal boundaries and areal subdivisions.

Because the EVL measures are potentially influenced by variation in cortical thickness, we also compared the EVL and cortical thickness gradients (Fig. 4C, D). The modality specific gradients largely differed in location, indicating that these layer-specific T1w/T2w-FLAIR gradient-ridges were not simply attributable to the variations in cortical thickness.

Comparison with previous macaque atlases, generated using ex vivo histology, is shown in Figure 5 (Van Essen et al., 2012). While there is evident convergence among the atlases in the unimodal sensory cortices, as well as transmodal cortices such as intraparietal cortex (Lewis and Van Essen 2000), there are more notable differences in the higher level cognitive regions such as prefrontal and orbitofrontal cortices as well as early peri-allocortical regions. These differences likely reflect multiple factors including different methodological strengths and weaknesses, species (e.g. mulatta vs cynomolgus) and sex (male vs female) and potential subject variability across the studies.

**Figure 5.**
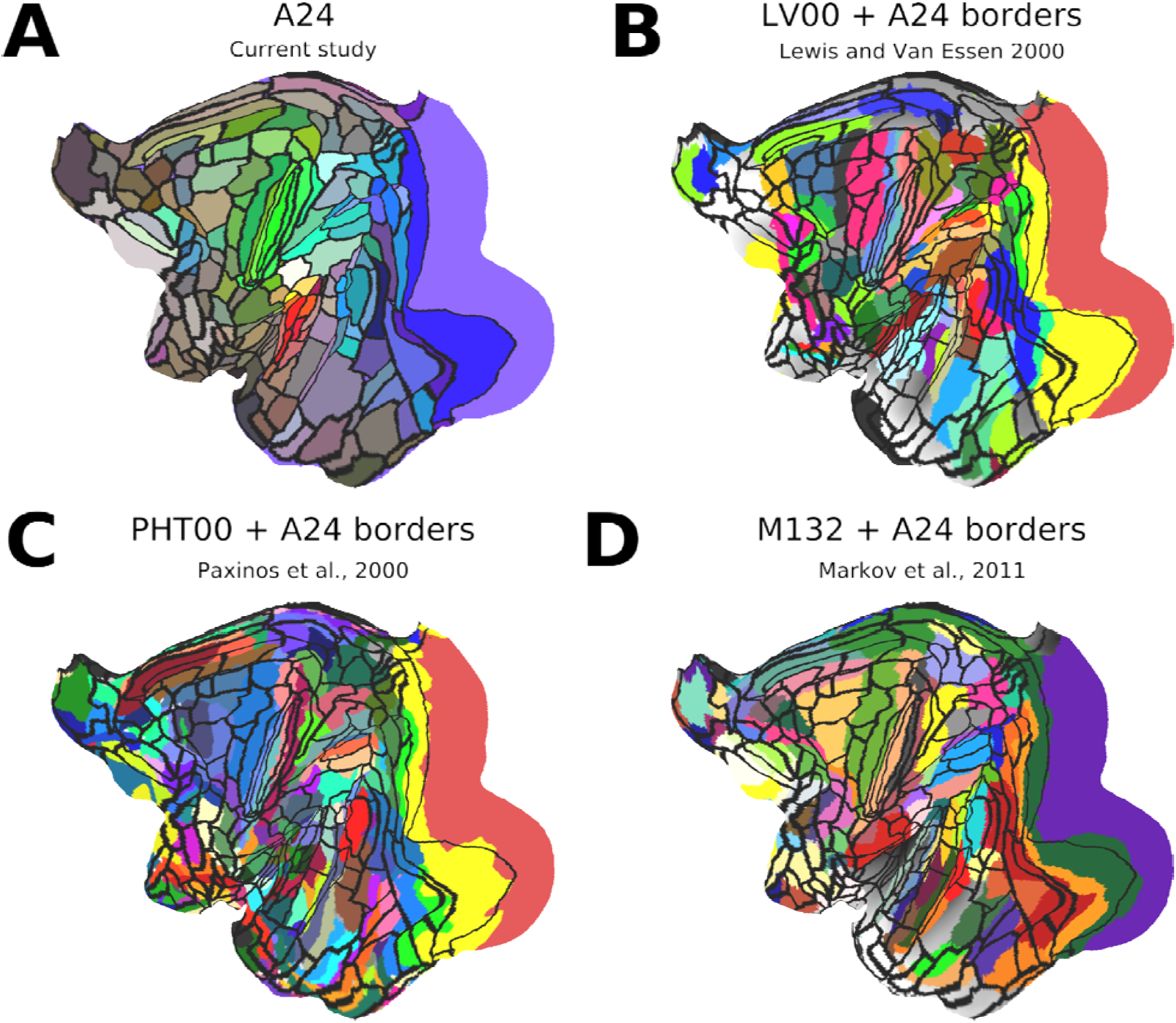
Comparison of Cortical Area Atlases from Four Studies. **(A)** Cortical area boundaries estimated in the current study using layer-specific myeloarchitectonic transitions. **(B)** Lewis and Van Essen (LV00) multi-modal histology atlas (Lewis and Van Essen 2000). **(C)** Paxinos, Huang, and Toga (PHT00) multi-modal histology atlas (Paxinos et al., 2000). **(D)** M132 cytoarchitecture atlas (Markov et al., 2011). Each atlas is overlaid with borders (black contours) estimated in the current study. The LV00, PHT00 and M132 atlases were compiled by Van Essen and colleagues (Van Essen et al., 2012) and obtained from the BALSA repository (Van Essen et al., 2017). Note that different macaque species were used in the studies: A24 macaque mulatta (rhesus); LV00 macaque fascicularis (cynomolgus); PHT00: macaque mulatta and M132: macaque fascicularis. Data at https://balsa.wustl.edu/TBA.

Cortical Layer Myeloarchitecture Links with Hierarchical Organization of the Cortex.To investigate systematic variations in cortical myeloarchitecture, we applied agglomerative clustering on parcellated group-concatenated T1w/T2w-FLAIR EVLs (Fig. 6B, C, D). The dendrogram (Fig. 6C) revealed four distinct cortical types across sensory and association areas, primarily determined by the myelin density (Fig. 6A). This clustering revealed a myeloarchitectonic progression from agranular paleocortical, and surrounding dysgranular areas toward eulaminate cortices (Fig. 6E) (Barbas 1986, Garcia-Cabezas et al., 2020, John et al., 2022). The average cluster profiles showed most notable myelin differences in the middle cortical layers, with uni- and transmodal areas exhibiting relatively higher myelin content than found in supramodal association areas (Fig. 6F). These differences may be attributed to the inter-areal and cortico-thalamo-cortical connection differences between granular and agranular cortex (Fig. 6H).

**Figure 6.**
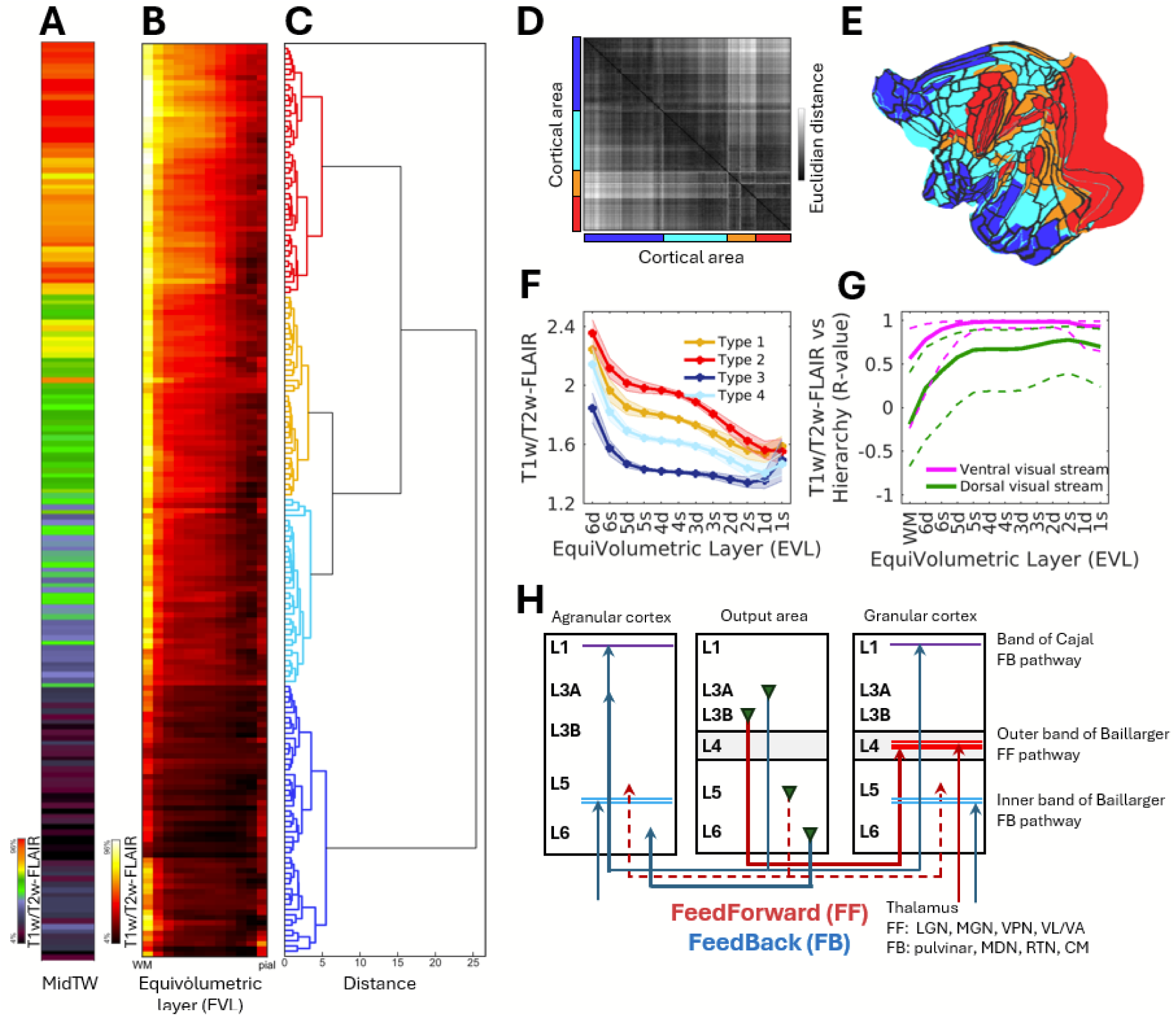
Laminar Myeloarchitecture Intricately Links with Cortical Hierarchy. **(A)** Midthickness-weighted and **(B)** equivolumetric layer (EVL) T1w/T2w-FLAIR myelin ordered according to **(C)** dendrogram. **(D)** Euclidean distance similarity matrix. **(E)** Clusters distribution displayed on a flat-map and **(F)** their average laminar profiles. Error-bar indicates standard deviation across cluster parcels. **(G)** T1w/T2w-FLAIR equivolumetric layers were highly correlated with the hierarchy levels of the ventral and dorsal visual streams. Hierarchy levels were determined using feedforward and feedback connections and obtained from the literature (Van Essen et al., 1991). **(H)** Integrated model of dual counter-stream architecture and bands of Baillarger (Markov et al., 2013). Feedforward (FF; red lines) and feedback (FB; blue lines) projections. FF and FB pathways, as well as the inner and outer bands of Baillarger, receive substantial contributions from distinct thalamic nuclei (Rockland et al., 1989, Jones 2001, Hubel et al., 1972). Abbreviations: CM: Centromedial Nucleus; LGN: Lateral Geniculate Nucleus; MDN: MedioDorsal Nucleus; MGN: Medial Geniculate Nucleus; RTN: Reticular Nucleus; VL/VA: Ventral Lateral/Ventral Anterior Nucleus; VPN: Ventral Posterior Nucleus.

Given the similarity between laminar myelination and cortical types, we further corroborated the link between layer myeloarchitecture and areal microcircuit specialisation (Felleman and Van Essen 1991, Burt et al., 2018). For this objective, we correlated the hierarchical level of areas in ventral and dorsal visual streams, determined by feedforward and feedback connections (Felleman and Van Essen 1991), with cortical myelination across the EVLs. We found that significant correlation between hierarchy level and myelination extended across all EVLs, except in EVL6d adjacent to WM (Fig. 6G).

## Discussion

Using cortical layer thickness-adjusted, Gibbs’ ringing attenuated T1w/T2w-FLAIR MRI, we present a methodology for charting cortical areas based on their distinctive laminar myeloarchitecture. Comparisons between in vivo myelin measures and ex-vivo myelin histology revealed strikingly similar layer profiles and sharp, layer-specific gradient-ridges revealing cortical area boundaries substantiating the validity of the methodology. Altogether, these methodological and conceptual advances enable precision myeloarchitectonic imaging across cortical areas and species, and represent a significant advance in noninvasive parcellation of the cerebral cortex.

T1w/T2w mapping was developed as an accessible, high CNR non-invasive measure of cortical myelination (Glasser and Van Essen 2011). The specificity of this approach for myelin, however, has been criticised, with many alternative, more quantitative approaches being proposed. These alternatives often face limitations in achievable resolution, signal-to-noise ratio, scan time, B_0_ field strength requirements, and availability of pulse sequences (Glasser et al., 2022). Here, we addressed this criticism by demonstrating that using T1w/T2w-FLAIR, up to 90% of the variance is attributable to myelin (Fig. 2). These results are in good agreement with postmortem study demonstrating that T1w is strongly correlated with myelin (Eickhoff et al., 2005, Leuze et al., 2017), but extend such measures to whole-cortex layer mapping in the living subjects by capitalising on sequences that provide anatomically accurate data needed for precise reconstruction of the equivolumetric surfaces (Supp. Fig. 2).

Charting the cortical layers enables exploring more specific neural populations (pathways), enhancing our understanding of neuroanatomy but also contributes to issues concerning disease, development, and evolution. For example, our results revealed the division of M1 into caudal and rostral sections (Fig. 3), likely corresponding to the evolutionary “new and old M1’’’, respectively (Rathelot et al., 2009). In M1c, cortico-motoneuronal (CM) cells make connections with motorneurons innervating muscles and allow for highly skilled movements and are only present in higher primates, including humans. Our results suggest that such anatomical division is also present in the nonhuman primate face region (Fig. 3A, B), which may support skilled use of the mouth and tongue, essential components for the emergence of voice articulation and language in humans, but see (Cerkevich et al., 2022). Indeed, primary lateral sclerosis (PLS) is a neuromuscular disorder characterised primarily due to the degeneration of upper motor neurons in the brain, which are CM cells, and is associated with dysarthria (slurred speech). This contrasts with amyotrophic lateral sclerosis (ALS), where both upper and lower motor neurons degenerate often leading to anarthria (complete loss of speech). Viewed from a clinical perspective, given the human median cortical thickness is 2.7 mm (Hayashi et al., 2022), scaling the image resolution to standard six cortical layer-model corresponds to about 450 µm isotropic. This implies that these clinically relevant anatomical features may be within reach of in vivo human MRI applications when combined with emerging 3D EPI (Stirnberg et al., 2022) and image de-noising technologies (Vizioli et al., 2021).

The progressive reduction in the layer myelination from unimodal primary areas towards supramodal association areas (Fig. 6B), likely reflects variations in fundamental circuit motifs (e.g. FF/FB and thalamocortical pathways, and column/module surface area) along the hierarchical organisation of the cerebral cortex (Felleman and Van Essen 1991, Markov et al., 2013, Burt et al. 2018) (Fig. 6G, H). Feedforward connections project from supragranular layers to granular and superficial layers (dense L3B→L4 and sparse L5→L5), whereas feedback projections project from infragranular layers to infragranular layers and cortical layer I (sparse L5→L5 and dense L6→L1). This implies that in the granular cortex the myelin in middle cortical layers are more weighted by the feedforward axons. In contrast, dysgranular and agranular cortex, characterised by diffuse and absence of L4, myelin is more weighted by the feedback axons. Moreover, myelin’s inhibitory effect on synaptic connections (Pajevic et al., 2014, Glasser et al 2015, Froudist-Walsh et al., 2024) may be critical to prevent mixing feedforward and feedback circuits thereby enabling structurally distinct and functionally meaningful dual-counterstream architecture enabling comparison between internally generated predictions and external sensory data (Friston 2010, Bastos et al., 2012, Vezoli et al., 2022).

Additionally, myelinated thalamo-cortical axons, situated in the outer and inner bands of Baillarger, relative to the canonical circuitry (Douglas et al., 2007), are positioned after the primary output layers of inter-areal feedforward and feedback projections, respectively (Fig. 7H). This suggests that structurally the outer band of Baillarger is part of the feedforward pathway whereas the inner band of Baillarger is part of the feedback pathway. Such dichotomy would allow the thalamus to regulate neural synchronisation and system-level information processing by narrowing neural time delays independently in feedforward and feedback channels (Salami et al., 2003, Pajevic et al., 2014). Overall, this model integrating dual-counterstream architecture and bands of Baillarger may provide new explanatory power for cortical laminar myeloarchitecture (Fig. 6B, G, H).

**Figure 7.**
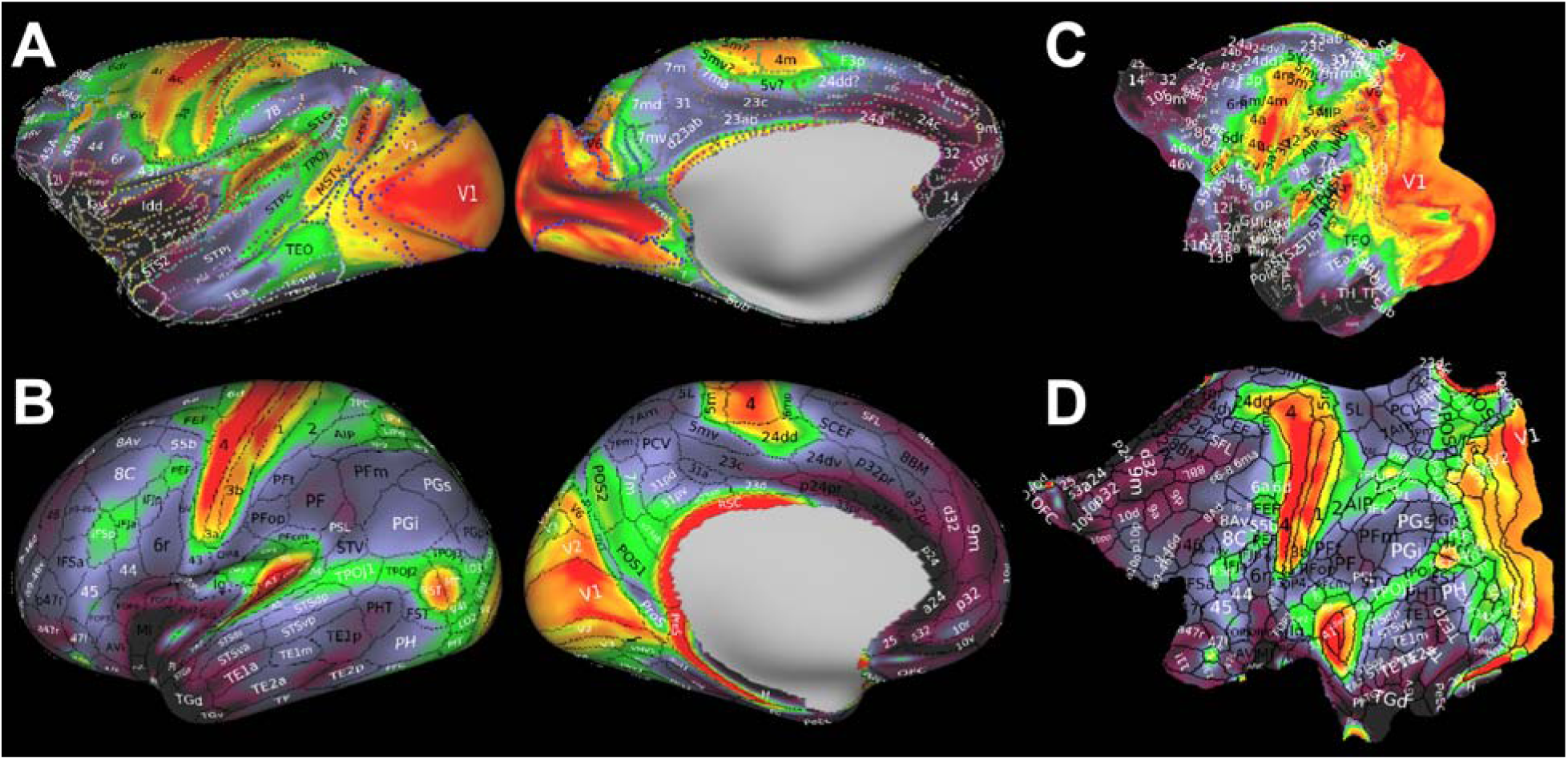
Comparison of macaque and human parcellation of cerebral cortex. **(A)** Macaque 182-area boundaries delineated using T1w/T2w-FLAIR laminar myelin MRI. **(B)** The human connectome project 180-area parcellation delineated using multi-modal MRI (Glasser et al., 2016). **(C, D)** Parcellations displayed on cortical flat-maps. Parcellations are overlaid on midthickness-weighted myelin maps. Putative homologies beyond primary sensory areas can be identified by myelin and topographic organisation between cortical areas. Data at https://balsa.wustl.edu/TBA.

However, we found that the association between cortical myelination and hierarchical organisation was unexpectedly strong also in the superficial EVLs (Fig. 7G). While in the middle cortical layers this association may, in part, reflect feedforward and feedback pathways and bands of Baillarger (Fig. 7H), in the superficial layers the association may reflect increased cortical column/module surface area required for more complex neural presentations at the higher levels of cortical hierarchy (Elston 2007).

The layer-mapping of cortical myeloarchitecture provides a powerful addition to our multi-modal comparative imaging battery for studying the organisation of primate brains (Markov et al., 2013, Glasser et al., 2016, Autio, et al, 2020, Hayashi et al., 2021). Layer-specific myeloarchitectonic transitions and putative homologous area identities between macaque and human offer improved landmarks for interspecies registration (Fig. 7), potentially improving our understanding of the evolutionary expansion of the cerebral cortex (Van Essen 2007). Our results exhibit a substantially larger range of myelin gradients than previously reported (Glasser et al., 2016, Donahue et al., 2018), which may enable suggestions for new putative homologues between macaque and human using myelin density and topological arrangement of the areas.

However, it is challenging to determine area boundaries using myelin alone. For instance, in the visual system, myeloarchitectonic variations may partially reflect different circuit properties related to central vs peripheral vision (Tootell et al., 1982). In V1, the central visual representation exhibits relatively sparser myelination compared to the V1 peripheral presentation, potentially due to differences in column/module and neuron densities (Tootell et al., 1982, Collins et al., 2010) and connectivity (Wang et al., 2022), in both macaque and human (Fig. 7C, D). Such retinotopic and myelin relations may be present in other visual areas as well, leading to erroneous boundary estimations. Similar challenges are met along the sensori-motor homunculus (Fig. 3A, B; Supp. Fig. 4). Therefore, it is essential to consider these myelin gradients in the context of functional imaging and multi-modal histology to refine the presented area boundaries and their identities (Glasser et al., 2016, Van Essen et al., 2018, Hayashi et al., 2021, Avram et al., 2021).

Notwithstanding, the remarkable similarity between macaque and human cortical myeloarchitecture is intriguing and warrants further comments (Fig. 7). For instance, in the medial parietal cortex, we observed an intermediate stripe of myelin in both species (Fig. 7A, B). However, in macaques, this region suggested to be part of the default-mode network (Yokoyama et al., 2021) or V6 complex (Galletti et al., 1999) while in humans, it contains higher-visual processing areas (e.g., POS1 and POS2) that are likely human-specific (Glasser et al., 2016). One possible explanation for this dilemma is that inter-areal and cortico-thalamo-cortical connections may have undergone distinct evolutionary selection processes. The density of inter-areal connectivity is known to reduce with increasing brain size, resulting in structurally sparser and functionally more independent areas (Ventura-Antunes et al., 2013). In contrast, the size of the thalamus is directly proportional to the number of neurons in the cerebral cortex (Herculano-Houzel et al., 2008), suggesting that the density of myelinated cortico-thalamo-cortical axons may be relatively preserved across primate evolution. Therefore, the striking similarity between macaque and human cortical myeloarchitecture may reflect a shared developmental program of cortico-thalamo-cortical connections (Lopez-Penditto et al., 2003), while the increased number of functionally independent areas may indicate species-specific sparsity of inter-areal connectivity and enlarged receptive fields necessary for the higher-order functions of the human brain.

## Conclusions

Here, we presented a novel cortical layer thickness-adjusted Gibbs’ ringing attenuated T1w/T2w-FLAIR methodology to chart layer-specific myeloarchitectonic transitions in primate cerebral cortex. Our results demonstrate that cortical-layer specific area boundaries, and area-subdivisions, can be noninvasively mapped in living primates, thereby advancing the pioneering postmortem work of classical neuroanatomists towards an era of non-invasive comparative cortical layer myelin imaging.

## Methods and Materials

### Data Acquisition

Experiments were performed using a 3T MRI scanner (MAGNETOM Prisma, Siemens Healthcare AG, Erlangen, Germany) equipped with 80mT/m gradients (XR 80/200 gradient system with slew rate 200T/m/s), a 2-channel B_1_ transmit array (TimTX TrueForm) and a 24-channel coil for macaque brain (Rogue Research Inc, Montreal/Takashima Seisakusho, K.K., Tokyo) (Autio et al., 2020). The animal experiments were conducted in accordance with the institutional guidelines for animal experiments, and animals were maintained and handled in accordance with "Guidelines for the Care and Use of Nonhuman Primates in Neuroscience Research" by the Japan Neuroscience Society, and “Guidelines for the Care and Use of Laboratory Animals” (National Institute of Health, Bethesda, MD). All animal procedures were approved by the Animal Care and Use Committee of the Kobe Institute of RIKEN (MA2008-03-19).

### Anaesthesia Protocol

Macaque monkeys (Macaca Mulatta, N=7, mean=6.9 ± 1.0 kg, age=8.2 ± 1.9 years) were initially treated with atropine sulphate (20 μg/kg) and then sedated with intramuscular injection of dexmedetomidine (4.5 µg/kg) and ketamine (6 mg/kg). A catheter was inserted into the caudal artery for blood-gas sampling, and endotracheal intubation was performed for steady controlled ventilation using an anaesthetic ventilator (Apollo, Drager, Germany). End-tidal carbon dioxide was monitored and used to adjust ventilation rate (0.2-0.3 Hz) and end-tidal volume. After the animal was fixed in an animal holder, deep anaesthesia was maintained using 1.2–1.5% isoflurane via a calibrated vaporizer with a mixture of air 0.75 L/min and O_2_ 0.1 L/min. Animals were warmed with a blanket and their rectal temperature (1030, SA Instruments Inc., NY, USA), peripheral oxygen saturation and heart rate (7500FO, NONIN Medical Inc., MN, USA) were monitored throughout the experiments.

### Structural MRI Acquisition Protocol

T1w images were acquired using a 3D Magnetization Prepared Rapid Acquisition Gradient Echo (MPRAGE) sequence (320 µm isotropic, FOV=123×123×82 mm, matrix=384×384, slices per slab=256, sagittal orientation, readout direction FH, phase direction AP, averages=12-15, TR=2200 ms, TE=3.0 ms, TI=900 ms, flip-angle=8°, bandwidth=200 Hz/pixel, no fat suppression, GRAPPA=2, reference lines PE=32, turbo factor=224, averages=12-15, and pre-scan normalisation). T2w images were acquired using a Sampling Perfection with Application optimised Contrast using different angle Evolutions (SPACE)-FLAIR sequence (320 µm isotropic, FOV=123×123×82 mm, matrix=384×384, slice per slab=256, sagittal orientation, readout direction FH, phase direction AP, TR=5000ms, TE=397 ms, TI=1.8s, bandwidth=420 Hz/pixel, no fat suppression, GRAPPA=2, reference lines PE=32, turbo factor=188, echo train duration=933 ms, averages=6-7 and pre-scan normalisation). The total acquisition time for structural scans was ≈3 h.

### Myelin Histology

Surgical and histology procedures were in accordance with European requirements (86/609/EEC) and approved by the appropriate veterinary and ethical services. The experiment was conducted on a single Cynomolgus macaque (Macaca fascicularis, 15 year-old female, 5.2 kg). The detailed protocol for surgery, euthanasia and perfusion can be found elsewhere (Markov et al., 2014). The myelin stain was performed on mounted sections by adjusting the modified Gallyas silver stain method of which the details can be found in Pistorio et al, 2006. Briefly, after perfusion and extraction, the brain was cut in 40 μm-thick sections on a freezing microtome. 1 out of 24 sections (960 µm interval) were mounted on 3% gelatin-coated slices for myelin stain. After drying overnight, myelin silver staining begins with a fixation in formalin, followed by acetylation in 2:1 pyridine and acetic anhydride mixture and successive baths of decreasing concentration of pyridine. Those steps efficiently reduce the ability of the other tissue elements to retain staining, mainly cell nuclei, and ensure that myelin is the only compound that can effectively bind with silver particles. After rinsing with water, sections were bathed in silver nitrate solution for impregnation, the surplus of silver is then washed away by 0.5% acetic acid, followed by re-fixation in formalin. After the re-fixation, the sections are immersed in developer solution, thus revealing myelin fibres. This step is critical and must be visually monitored by experimentalists under a dissecting microscope. When the appropriate contrast is reached, the revelation reaction is stopped with 1% acetic acid, and histological contrast is adjusted by differentiation (i.e. destaining) of overdeveloped sections with 0.1% potassium ferricyanide solution. Sections are then washed by 0.5% sodium thiosulfate and water, dehydrated and coverslipped.

## Data analysis

### Structural MRI Processing

The data were preprocessed using the NHP version of the HCP Pipelines (Glasser et al., 2013, Autio et al., 2020). Each T1w and T2w-FLAIR image was corrected for gradient non-linearity and averaged in their mid-point space. Then, each T1w and T2w-FLAIR image was registered into an anterior-posterior commissural (AC-PC) alignment using a rigid body transformation, following B_0_ inhomogeneity distortion correction and brain extraction. T2w-FLAIR and T1w images were aligned to sub-voxel precision using boundary based registration (Greve and Fischl, 2009), and corrected for signal intensity inhomogeneity using B_1_- bias field estimate from the sqrt(T1w×T2w) method (Glasser and Van Essen 2011; Glasser et al., 2013). Next, data was transformed into a standard macaque space (Autio et al., 2024) by 12-parameter affine and nonlinear volume registration using FLIRT and FNIRT FSL tools (Jenkinson et al., 2002).

The cortical surfaces were constructed using the FreeSurferNHP pipeline using *FreeSurfer* v6.0 (Fischl, 2012). This process included scaling volumes, intensity correction, segmentation of the brain into cortex and subcortical structures, reconstruction of the white and pial surfaces and estimation of cortical folding maps and thickness. First, brain volumes data were scaled to twice as large as the original physical space. The intensity correction of the T1w volume was performed using FMRIB’s Automated Segmentation Tool (FAST) (Zhang et al., 2001). Subsequently, the intensity corrected T1w volume was segmented using a FreeSurfer subcortical segmentation tool optimised for macaque (Fischl et al., 2002; Autio et al., 2020). Additionally, the white matter segmentation was guided by a template for white matter skeleton to ensure the delineation of the thin white matter blades (Autio et al., 2020). Visual inspection of white matter segmentation was performed, with manual editing of aseg.mgz, wm.mgz and filled.mgz carried out as necessary (by JAA), to improve white surface estimation accuracy.

Following the white surface estimation, initial surface registration was performed using the animal’s white surface “sulc map” to the macaque surface template (Autio et al., 2020). The pial surface estimation initially utilised the intensity normalised T1w image and was further refined using the T2w-FLAIR image to aid in excluding dura and blood vessels (Glasser et al., 2013). Then, surface and volume data in scaled space were transformed back (‘rescaled’) into the animal’s physical space, and the cortical pial surface and thickness were estimated.

The PostFreeSurfer pipeline transformed anatomical volumes and cortical surfaces into the Yerkes19 standard space, performed a more optimised surface registration using folding information via MSMSulc (Robinson et al., 2018, 2014), and generated the mid-thickness surface (by averaging the white and pial surfaces), inflated and very inflated surfaces. The myelin map was generated from the T1w/T2w-FLAIR ratio and the volume to surface mapping of the T1w/T2w-FLAIR ratio was performed using a ‘myelin-style’ mapping (Glasser and Van Essen, 2011), in which a cortical ribbon mask and a metric of cortical thickness were used, weighting voxels closer to the midthickness surface. Voxel weighting was done with a Gaussian kernel of 3 mm FWHM. The surface models used for mapping the T1w/T2w-FLAIR ratio were the FreeSurfer native mesh, and all the surface geometry and metric data were resampled to a high-resolution 164k standard mesh (per hemisphere) that had intrinsic left/right hemisphere correspondence of vertices (Van Essen et al., 2012) using the MSMSulc surface registration (Robinson et al., 2014, 2018, Autio et al., 2020).

High-resolution rhesus (N=7; RhesusHighRes7) group average surfaces, curvature, cortical thickness and cortical midthickness-weighted T1w/T2w-FLAIR maps were generated using the HCP-NHP MakeAverageDataset pipeline. Then, these maps were minimally smoothed along the cortical surface (300 µm), and gradient maps were calculated using wb_commands -cifti-smooth and -cifti-gradient, respectively.

### Equivolumetric Surface Mapping

Twelve cortical layer surfaces were generated based on equivolume model (Bok 1929, Waehnert et al. 2014; Wagstyl et al., 2018, Consolini et al., 2022) using the Workbench command ‘-surface-cortex-layer’ and the native pial and white surfaces in the subjects’ physical AC-PC aligned space. The original T1w and T2w-FLAIR 0.32 mm images were transformed to the subject’s AC-PC space, corrected for B_0_ distortion and gradient nonlinearity, averaged, interpolated with cubic splines to 250 µm resolution and then mapped to native laminar mesh surfaces using Workbench command ‘-volume-to-surface-mapping’ with the ribbon-constrained algorithm.

A rigorous quality assurance process was conducted for the EVLs near the white matter and pial surfaces. When errors in white matter surface placements were identified, manual editing of aseg.mgz, WM.mgz and filled.mgz files were performed. If the pial surface extended to a vessel running along it (pial vasculature is readily visible in high-resolution T1w images), the surface was repositioned using FreeSurfer 7.0 and Freeview 3.0 (by JAA). After these corrections, the FreeSurfer-NHP pipeline was re-run, and this process was repeated until the errors in the volume space became visually negligible.

Since myelinated axons traverse parallel to the normal of the cortical surface, and bands of Baillarger are oriented normal to the cortical surface, the orientation of the cerebral cortex relative to the direction of B_0_ may bias signal intensity of T1w and T2w-FLAIR images differentially and subsequently the T1w/T2w-FLAIR ratio (Wharton et al., 2012, Bartels et al., 2022). We assume that B_0_ orientation bias ∝ cos^2^(ϴ), where Θ is the angle between normal of the cortex relative to the direction of B_0_. Each vertices ϴ was determined in the subject’s original MRI space (Autio et al., 2024). Then, orientation bias and cortical curvature were regressed out from each contrast and each EVL surface map.

Applying MSMSulc surface registration, the data was resampled to the atlas space using ADAP_BARY_AREA with vertex area correction, and left and right hemispheres were joined into a CIFTI file.

### Charting Cortical Area Boundaries

To investigate sharp, cortical-layer specific myeloarchitectonic transitions, group-average B_0_ orientation and curvature corrected T1w/T2w-FLAIR EVL maps were smoothed using a factor of 300 µm and spatial gradient-ridges were calculated using wb_command -cifti-gradient. When spatial myelin gradients displayed symmetry across the hemispheres and did not explicitly follow cortical curvature, putative area/subarea boundaries were manually drawn using the wb_view borders function (by JAA). The resulting border.gii file, containing the area boundaries, was converted to metric format using wb_command -border-to-rois and then to dlabel format using wb_command -cifti-create-label.

To investigate repetitive patterns in cortical layer myeloarchitecture, the T1w/T2w-FLAIR EVL profiles were parcellated using the estimated area borders. Because some of the outermost parcellated T1w/T2w-FLAIR EVLs exhibited inhomogeneity due to artefacts (e.g., FreeSurfer errors in pial surface estimation, areas adjacent to major vessels), median values were assigned to each parcel in each EVL. Subject data were then concatenated (N=7, and an additional test-retest data set), EVLs and hemispheres (8×12×2=192). Then agglomerative clustering was performed using Ward’s method. A dendrogram was used to determine the optimal number of myeloarchitectonic clusters.

### Comparison of Laminar Myelin Stain and MRI Profiles

Modified Gallyas myelin sections were imported to Zen software (3.5 Blue Edition, Carl Zeiss Microscopy GmbH, Oberkochen, Germany) and ROIs were drawn perpendicular to the cortical surface. Corresponding ROIs were determined using MRI volume data, drawn on the midthickness weighted cortical surface and the areal identity was confirmed using the M132 areal atlas (Markov et al., 2014). The ROIs (stain and MRI) were then imported to Matlab, the stain was averaged in the radial direction and down-sampled to 12 data points in the layer direction, and maximal correlations were searched to overcome white matter surface placement errors (in both myelin stain and MRI) using a 10 μm sliding-window bounded within ±150 μm from the observer-determined WM surface.

## Author Contributions

Conceptualization: JAA; Methodology: JAA; Software: JAA AU KM TC TH; Formal analysis: JAA AU TI IK TH; Investigation: JAA TO LM YH MO YM; writing - original draft: JAA; writing - review & editing: JAA TI LM HK DVCE MFG TH; visualisation: JAA TI KM; project administration: TH, TO; Funding acquisition: JAA HK DVCE MFG TH.

## Acknowledgements

This research is partially supported by the program for Brain/MINDS and Brain/MINDS-beyond from Japan Agency for Medical Research and development, AMED (JP18dm0307006, JP21dm0525006, JP19dm0307004, JP23wm0625001, JP24wm0625205, JP24wm0625122, T.H.), NIH BICAN project (UM1MH130981, T.H., D.V.E., M.F.G., H.K.), JSPS KAKENHI (JP22H04926, JP23H00413, T.H.; JP20K15945, J.A.A.; JP15K08707, T.O.), NIH R01 MH60974 (D.V.E., M.F.G.), NIH T32 EB021955. French National Research Agency (LABEX CORTEX ANR-11-LABX-0042. H.K.; Université de Lyon ANR- 11-IDEX-0007. H.K.; A2P2MC ANR-17-NEUC-0004, CORTICITY ANR-17- HBPR-0003, H.K.), SCUSI, Région Auvergne-Rhône-Alpes (1700933701, H.K.), Chinese Academy of Sciences President’s International Fellowship Initiative (Grant No. 2018VBA0011, H.K.), Brain research Foundation (FRC APE13 “Rotary-Espoir en Tête”, H.K.). The authors appreciate technical contributions and discussions from Reiko Kobayashi, Yuki Matsumoto, and Yuta Urushibata.

