## Supplemental Figure 1-8 for "Charting cortical-layer specific area boundaries using Gibbs’ ringing attenuated T1w/T2w-FLAIR myelin MRI"

**Affiliations**

Incl. **Supplementary Figures S1-9** and supplementary discussion.

**Data Acquisition Strategy - Attenuating Gibbs’ Ringing Artefacts**

Laminar myeloarchitectonic MRI has the potential to be a valuable tool in neuroanatomy and radiology. However, several challenges need to be addressed to obtain neuroanatomically meaningful laminar profiles. These challenges include image acquisition artefacts (e.g., B_1_ transmission and receive biases, Gibbs’ ringing, and B_0_ orientation bias) and data analysis biases (e.g., image co-registration, precise cortical surface reconstruction, and removing blood vessels) (Glasser et al., 2011; 2022; Autio et al., 2024). Here, we focus on describing the Gibbs’ ringing artefact and its impact on genuine cortical layer imaging contrasts.

The highly convoluted geometry of the primate cerebral cortices imposes a major challenge to MR image reconstruction. It is well known that the Fourier transformation of the complex MR signal (sine waves) at a sharp signal transition (e.g. box-function) such as at the CSF-cortex interface results in Gibbs’ ringing artefact (Fig. 1A, B). This can be detrimental to laminar analysis of MRI because the ringing artefacts manifest orthogonal to the cortical layers (Fig. 1B, inset) it can be confounded with genuine anatomical layers. Moreover, multiple sharp signal-intensity edges (e.g. opposing banks of sulci in central sulcus) generate superimposed sinc-waves that may form constructive (in-phase sinc-wave) or destructive (out-of-phase sinc-wave) interferences that increase or decrease ringing artefacts, respectively. The magnitude of the Gibbs’ overshoot also depends on the precise location of the edge relative to the image sampling grid (Ferreira et al., 2009; Kellner et al., 2016).

Because the image contrast differs between T1w and T2w images, Gibbs’ ringing artefacts also differ. In the T1w image, the signal intensity in CSF is very low in comparison to CGM and the Fourier expansion of a CSF-CGM interface causes an oscillation (‘Gibbs’ ringing’) with a signal overshoot in the outermost voxel in the CGM (Fig. 1A,B). In contrast, the signal intensity in the T2w image is relatively higher in the CSF compared to CGM and the Fourier transformation at the CSF-CGM interface causes a ringing artefact with a signal undershoot in the outermost voxel in the CGM (Fig. 1A,B). Because T1w and T2w images are contaminated by ringing artefacts (e.g. sinc waves) in opposing phases, the T1w/T2w ratio amplifies the ringing artefact (Fig. 1C).

The Gibbs’ ringing artefact can be confirmed by altering the spatial resolution, which changes the spatial frequency of the oscillation (Supp. Fig. 1A, B; red arrows). This Fourier transformation image reconstruction artefact occurs not only at the cranium/CSF interface but also at the CSF/CGM interface. The interaction and superimposition of ringing artefacts create a spatial periodicity of 2 voxels perpendicular to the cortical surface (Fig. 1B, Supp. Fig. 1A). The largest Gibbs’ artefact occurs when the CSF space is exactly 1 voxel (e.g anatomical high-frequency component), resulting in an in-phase Gibbs’ ringing artefact. The smallest artefact occurs when the CSF space is 2 voxels, resulting in an out-of-phase Gibbs’ ringing artefact. In addition, this superimposition of artefacts originating from opposing banks presents a further challenge for modelling. The ringing artefacts are prominent in the T1w/T2w (non-FLAIR) images (Supp. Fig. 1D, E; red arrows).

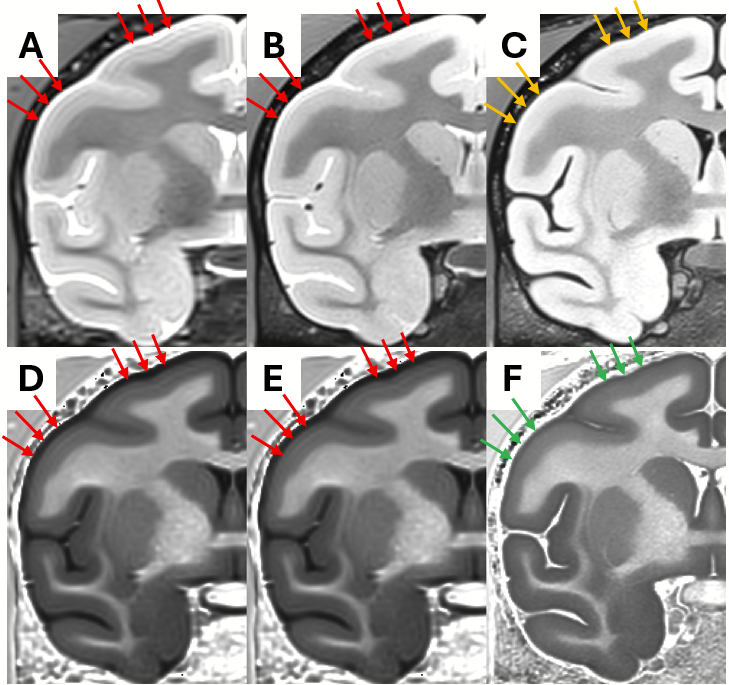

**Supplementary Figure 1. Gibbs’ Ringing Artefact Resembles Cortical Layers.** T2w-SPACE (wo-FLAIR) images acquired using **(A)** 500 µm and **(B)** 400 µm isotropic image resolution. Red arrows indicate prominent Gibbs’ ringing artefacts resembling cortical layers. Note that the frequency of the cortical signal-intensity oscillations are dependent on image resolution which is characteristic of Gibbs’ ringing artefact. **(C)** T2w-FLAIR exhibits notably smaller, yet still recognizable, Gibbs’ ringing artefacts (orange arrows) attributable to cerebrospinal fluid suppression. T1w/T2w (wo-FLAIR) myelin images created using **(D)** 500 and **(E)** 400 µm image resolutions. Note the prominent Gibbs’ ringing artefact in T1w/T2w images running beneath the pial surface (red arrows). **(F)** T1w/T2w-FLAIR does not exhibit notable Gibbs' ringing artefact (green arrows).

To overcome this limitation, we propose to calculate T1w/T2w-FLAIR ratio to simultaneously attenuate Gibbs’ ringing artefact (Fig. 1C) and to enhance the intracortical myelin CNR (Glasser et al., 2011). This approach is based on the idea that the *in-phase* ringing artefact can be selectively attenuated by dividing the two image types (here T1w and T2w-FLAIR) when their contrasts contain the same (spatial) high-frequency signal-intensity edges and polarity on the same discrete imaging grid, while also enhancing the contrast to laminar myelination.

The presented image ratio-based attenuation of the Gibbs’ ringing artefact is also applicable to other image-ratio-based contrasts, such as magnetization transfer ratio (MTR = 1 - S/S_0_), as long as the major ringing artefacts are *in-phase*.

An additional advantage of T2w-FLAIR is that the CSF suppression helps improve the image co-registration (T1w and T2w-FLAIR) and estimation of the pial surface (Supp. Fig. 2A). In the T2w (non-FLAIR) images it is challenging to precisely determine the signal intensity cut-off between CGM and CSF. In our experience, areas with high synaptic density tend to have underestimated pial surfaces because these regions exhibit notably higher signal-intensity in superficial layers of T2w images (data not shown). T2w-FLAIR overcomes this issue enabling precise delineation of white matter and pial surfaces that are essential for calculation of the EVLs (Supp. Fig. 2B).

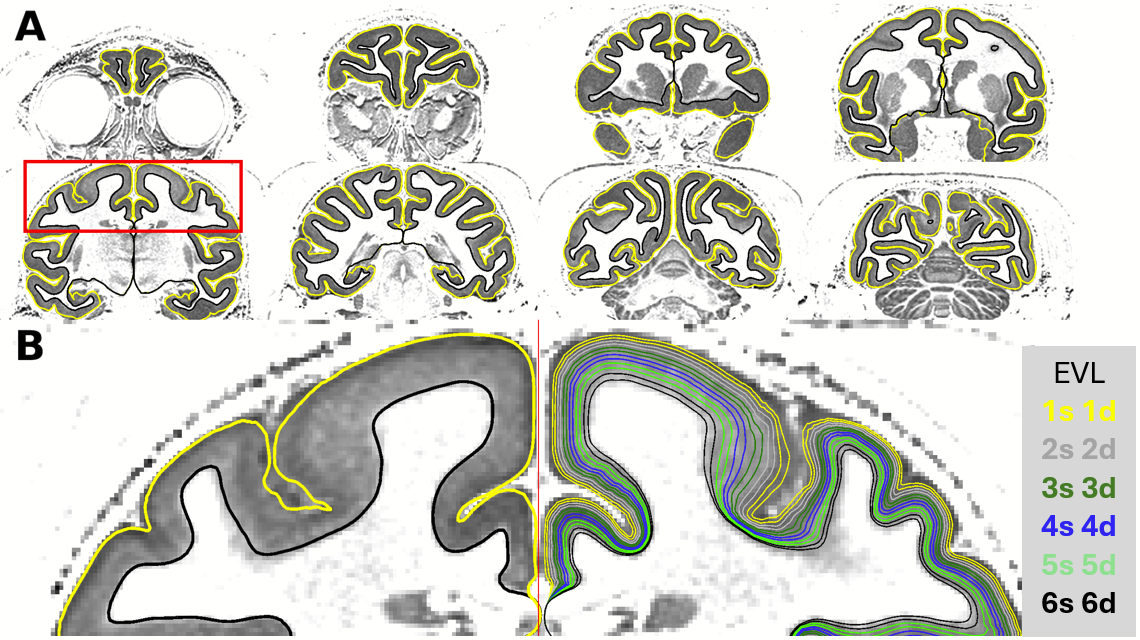

**Supplementary Figure 2.** **Cortical surface estimation in a representative macaque monkey.** **(A)** Pial (yellow) and white matter (black) surfaces were accurately delineated using a combination of high-resolution T1w and T2w-FLAIR images. Red rectangle indicates the location of the **(B**) zoomed view with equivolumetric layers (EVLs) displayed on the right hemisphere. Abbreviations: s: superficial; d: deep.

Increasing the image resolution also helps mitigate the effect of Gibbs’ ringing artefact (Supp. Fig. 1A, B) but it remains challenging within practical in vivo imaging resolutions that are close to the median layer thickness (assuming standard six-layer model). In terms of image resolution, achieving a target resolution that discerns sublayers, such as bands of Baillarger, is essential. The number of discernible sublayers varies and their thickness ranges from 40 to ≈500 µm, with some being diffuse (e.g. dysgranular cortex) and hardly distinguishable in histology (Von Economo and Koskinas, 1925). Nonetheless, our results imply that the minimum criteria for delineating cortical areas based on their layer-specific MRI contrasts is to scale the resolution to the approximate thickness of six standard cortical layers (Fig. 4) (Autio et al., 2021).

**Charting Bands of Baillarger**

While it was not our primary objective to make EVL maps of cortical sublayers, such as bands of Baillarger, it is noteworthy that the median cortical thickness-adjusted resolution allowed charting a large number of signal-intensity peaks throughout the cerebral cortex (Supp. Fig. 2). Detection of peaks, that are suggestive of Baillarger bands, was relatively robust in primary areas and adjoining multimodal areas (Fig. 2A, B). Unexpectedly, weak but systematic variations were observed in transmodal agranular cortices that lack or contain weak bands of Baillarger. For instance, the orbitofrontal cortex (11, 13, and 14) and parts of the insular cortex (e.g. areas lapl, lal, lai and lam) do not contain a notable band of Baillarger (Carmichael and Price 1994, Ervard et al, 2014). Thus, these peaks may have in part originated from other contrast mechanisms such as laminar variations in cell density (Fatterperker et al, 2001, Eickhoff et al., 2005), blood volume (Autio et al., 2024), or residual image processing artefacts (see Data Acquisition Strategy).

Regardless of the precise contrast mechanisms of these laminar signal-intensity variations, the peak distributions are highly suggestive of area transitions. For instance, (M1t vs area 3a) and subarea divisions (e.g. V1 central vision vs periphery).

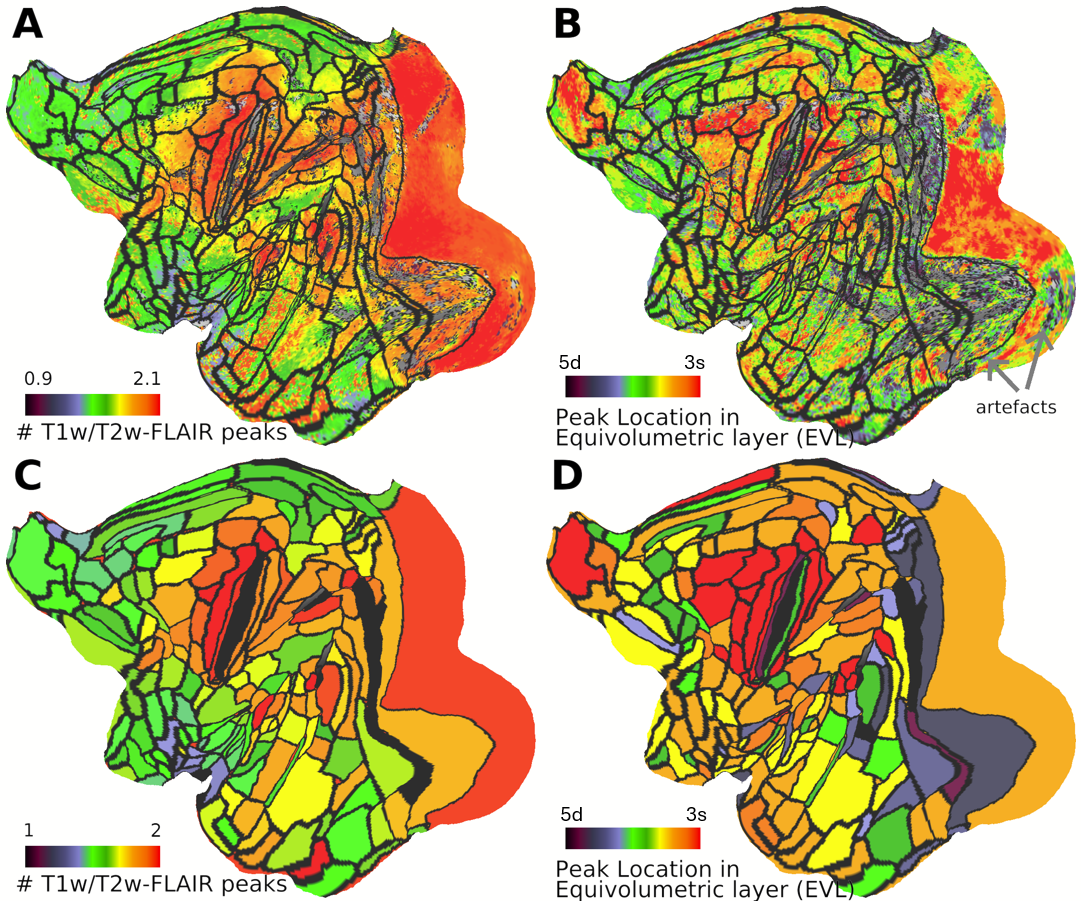

**Supplementary Figure 3. Charting Bands of Baillarger in the macaque cerebral cortex. (A)** Median number of T1w/T2w-FLAIR signal-intensity peaks (e.g. bands of Baillarger) across subjects. **(B)** Median location of T1w/T2w-FLAIR signal-intensity peak in equivolumetric layer (EVL). **(C, D)** Parcellated number peaks and peak locations.

**Somatosensory and Motor Cortex**

In the heavily myelinated sensorimotor cortices, several areas were delineated based on their distinct layer myeloarchitecture (Fig. 3). However, layer myelin imaging revealed unexpected complexity beyond the classical definition of somatosensory and motor cortices (M1, 3a, 3b, 2, 1 and 5) and further subdivisions along the somatotopy, making delineating this region challenging.

The primary motor cortex (M1), or area F1 (Matelli et al., 1985), was clearly distinguishable due to its strong myelin and thick cortex (Supp. Fig. 4G). However, myelin content in superficial EVLs varied within area F1 as defined by the M132 atlas (Supp. Fig. 4B,C, F). In EVL2b, myelin was high only in the caudal portion of the M1 (M1c). This region likely corresponds to Rathelot’s and Strick’s “new M1'', which contains cortico-motoneuronal (CM) cells in the more superficial layers (Rathelot et al., 2009). The rostral proportion of M1 (M1r), also known as “old M1'', does not exhibit myelin in EVL2, but in EVL3b and midthickness-weighted myelin maps, it shows notable higher myelination relative to the surrounding areas. Within M1r, there are notable gradients, and it remains unclear whether these correspond to motor homunculus or further subdivision of M1r. Myelin gradient-ridges, reminiscent of the motor homunculus, were also observed within M1c, and it will be interesting to compare these features with functional connectivity. The medial M1 (M1m) exhibits myelin homogeneity in EVL2b and has a slightly thinner cortex relative to M1c/r. Together, these results suggest subdivision of M1 aligning more with recent studies of connectivity (Rathelot et al., 2009) and receptor architecture (Rapan et al., 2021).

The boundary between M1c and area 3a revealed yet another conundrum: this zone exhibited high myelin (Supp. Fig. 2A), a band of Baillarger (Fig. 3C, Supp. Fig. 2A), sparse vascularity (Supp. Fig. 4H) and thick cortex relative to the bottom and caudal bank of the central sulcus. Although classically these features are not associated with area 3a, Krubitzer and colleagues have convincingly argued using electrophysiology and cytochrome oxidase histology that area 3a may extend to the caudal proportions of the sulcus (Krubitzer et al., 2004). On the other hand, it is widely recognized that area 3a can be determined by sparse myelin relative to the surrounding areas (Krubitzer et al., 2004; Burman et al., 2014; Glasser et al., 2016; Padberg et al., 2020). In our material, the relatively low myelin region corresponds to the bottom of the sulcus where the cortex is thin, which strongly suggests that area 3a is in the classically defined region. Another interpretation is that the M1c/3a zone indeed corresponds to area 3a (Krubitzer et al., 2004), and the additional myelin gradient in this region might correspond to the transition zone between areas 3a and 3b (Paxinos et al., 2000), or area 3a–hand–vestibular (3aNv) (Ödkvist et al., 1974, Guldin et al., 1998). Because area 3b does not have a well-developed outer band of Baillarger and M1 is characterised by a very wide myelination band, we refer to this zone as M1 transitory (M1t), albeit we acknowledge that the material presented herein does not allow us to unequivocally resolve this issue.

Relative to area 3a, area 3b exhibited denser myelin and vascularity and was delineated using the strong gradient-ridges (Autio et al., 2024). These results are consistent with findings from myelin (Fig. 3) and cytochrome oxidase stains (Krubitzer et al., 2004). However, myelin exhibits inhomogeneity within area 3b, likely corresponding to somatotopy: the upper limb area shows higher myelin in more superficial EVLs (Supp. Fig. 4B). The connectivity/cellular origin of this superficial myelin remains unclear, but a potential candidate is the area 3aNv (Ödkvist et al., 1974), reflecting the importance of vestibular-upper limb interaction for balance control in primates. Other theories propose that there may be additional subdivisions within M1 corresponding to motor commands and muscle synergies (Reilly et al., 2008) or effector-specific regions (foot, hand and mouth) and somato-cognitive action networks Gordon et al., 2023). Notably, the macaque tail region in 3a/3b exhibits a peculiar high myelination zone, and these zones approximately match with the action/body M1 subregions proposed by Gordon and colleagues (Gordon et al., 2023).

Sharp myelin transitions between architectonic areas 3b, 1, 2, 5d, 5v, 5m were clearly identifiable (Supp. Fig. 4D, E, I), despite these areas representing one of the thinner parts (1.5-2.0 mm) of the cerebral cortex (mean thickness 2.0 mm). In the medial sensorimotor cortex, strong myelin gradients suggested that areas 3a, 3b 2, and 1 do not extend as far medially as shown in M132 atlas (Supp. Fig. 4F). This view is also supported by abrupt reduction in vascularity (Supp. Fig. 2H, J) (Autio et al., 2024). The medial extent of sensorimotor cortex suggests topological homology with the HCP parcellation (Glasser et al., 2016). Finally, myelin inhomogeneity in areas 5m and 5d suggest that there may be more areas or somatotopy-related subdivisions (Pandya and Seltzer, 1982), and these variations deserve further investigations.

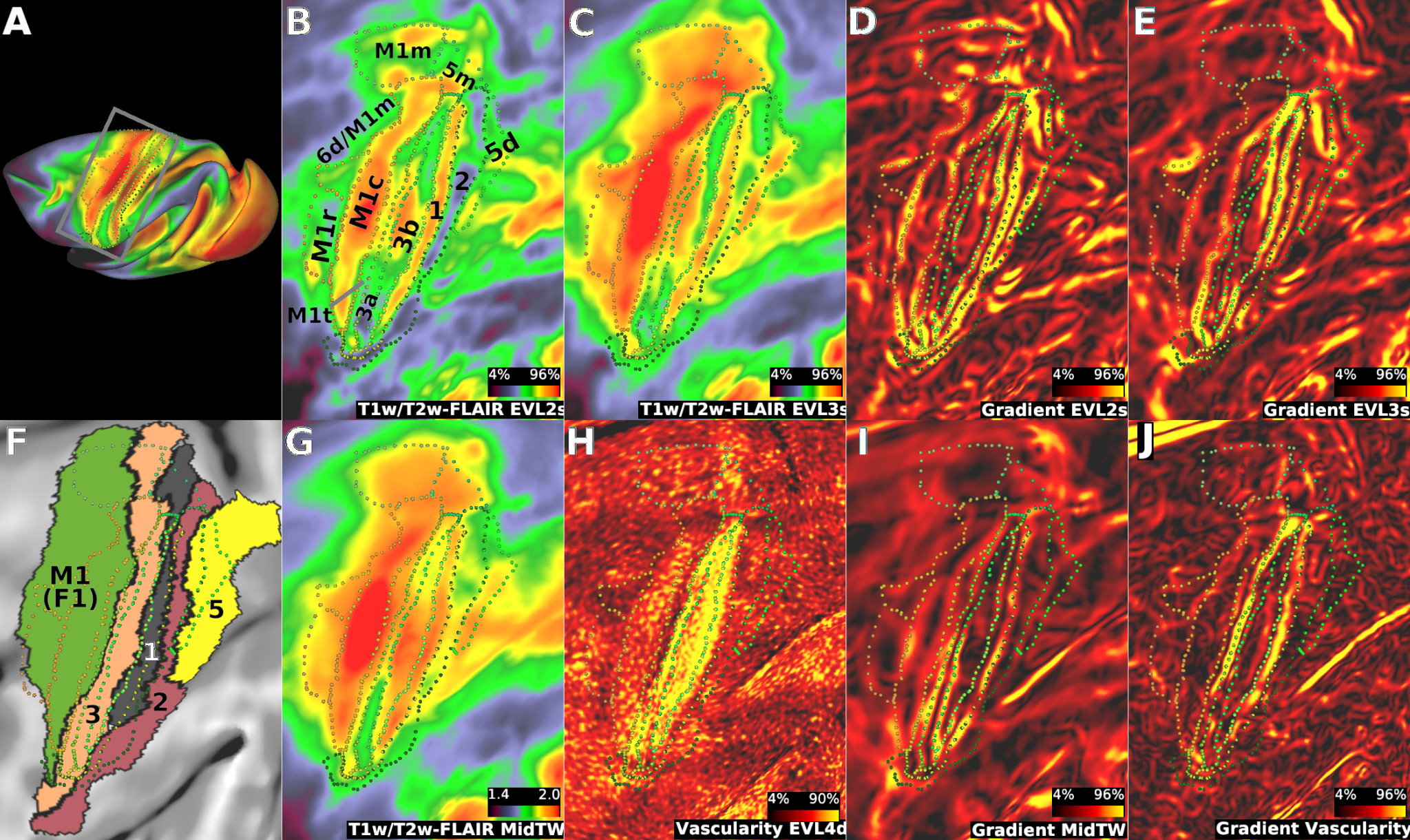

**Supplementary Figure 4. Charting Area Boundaries and Areal Subdivisions in the Sensorimotor Cortices. (A)** Location of the zoomed view (black box). **(B, C, G, H)** T1w/T2w-FLAIR from four representative equivolumetric layers (EVLs). **(F)** M132 atlas (Markov et al., 2011) overlaid with area boundaries of the current study. **(D, E, I, J)** Sharp gradient-ridges vary across EVLs, suggesting areal boundaries. Abbreviations: M1 (F1): Primary motor cortex; M1c: M1 caudal (Rathelot et al., 2009); M1r: M1 rostral (Rathelot et al., 2009); M1m: M1 medial (Rapan et al., 2021); M1t: M1 transitory (this study); 1: Brodmann area 1; 2: Brodmann area 2; 3a: Brodmann area 3a; 3b: Brodmann area 3b; 5d: Brodmann area 5 dorsal.

**Auditory and Vestibular Cortex**

In the auditory cortex, we identified a total of 13 areas (Supp. Fig. 5). Auditory cortex represents an exemplary region for demonstrating the advantage of mapping cortical layer myelin transitions, the advantage of averaging across layers for intermediately/sparsely myelinated cortex and the fine-grained complexity of the region.

The primary auditory cortex (A1), or core (Galaburda, Pandya 1983), was clearly distinguishable by dense myelin in the EVL3b (Supp. Fig. 5B). The rostral (R) and rostrotemporal (RT) subdivisions also exhibit strong myelin gradients (Hackett et al., 1998).

In the belt regions (Galaburda, Pandya 1983), we delineated five myelin gradient-ridges resembling the parcellation described by Hackett and colleagues (Hackett et al., 1998, 2011). We delineated three lateral (LBc, LBm, and LBr) and two medial (MBm and MBr) areas. The Hackett’s area MBc may very well exist within area 7op, however, due to the strong cortical curvature it was challenging to determine the location of the border.

In the parabelt region, we identified areas PBc and PBr, albeit the PBr was notably smaller in area than in previous parcellations (Hackett et al., 1999).

The complex subdivisions of the auditory cortex were recognized using the parcellation by Hackett and colleagues. Using myelin histology, they described a total of 13 areas: three core areas (A1, R and RT), four lateral belt areas (CL, ML, AL, RTL), four medial belt areas (CM, MM, RM and RTM), and two higher-level parabelt areas (PBc and PBr; Hackett et al., 1998, 2011). Overall, the laminar myelin progressively withdraws from superficial layers resembling the description of inter-areal connections in this region (Calaburda and Pandya 1983).

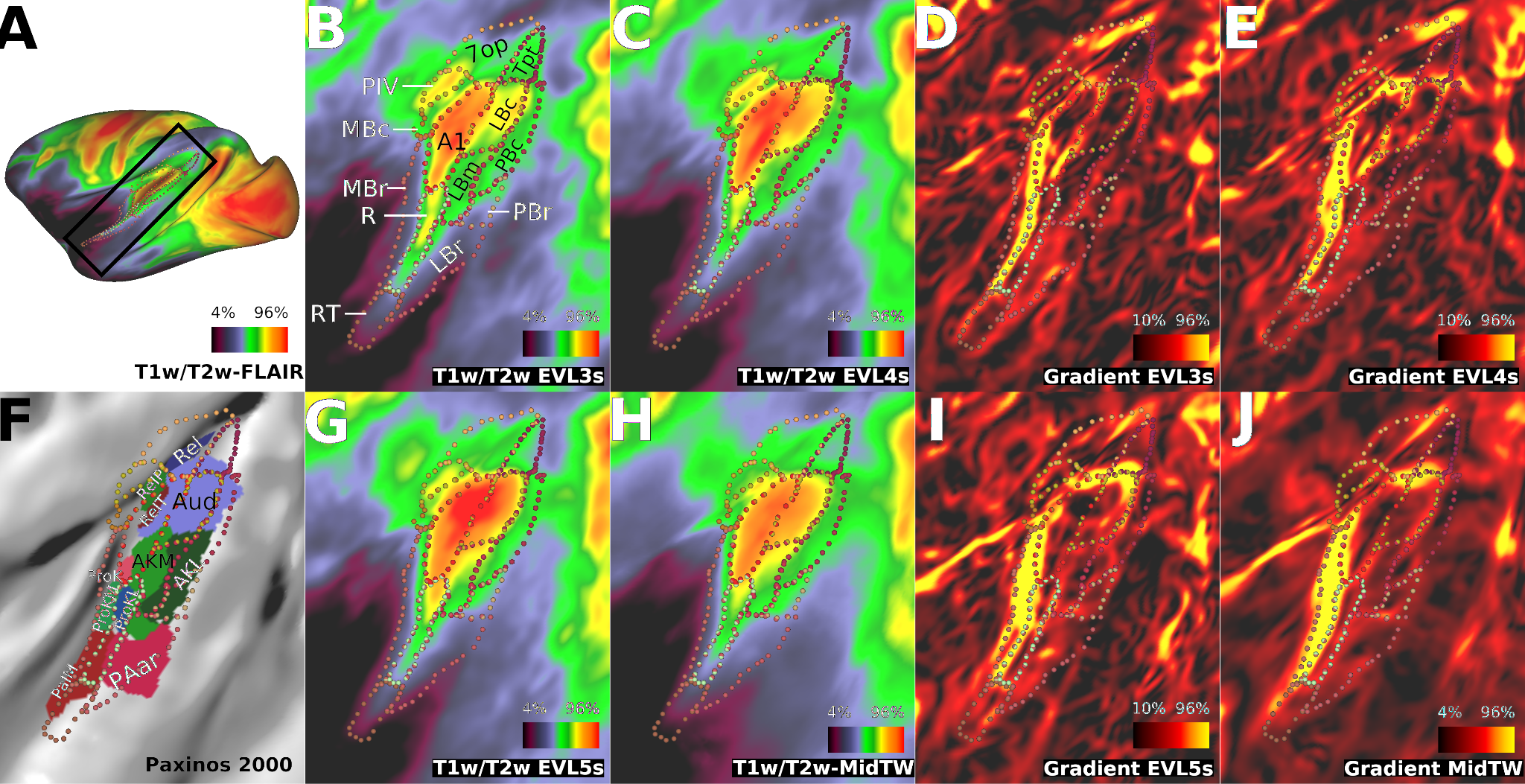

**Supplementary Figure 5. Charting Area Boundaries in the Auditory and Vestibular Cortex. (A)** Location of the zoomed view is indicated by the snippet (black box). **(B, C, G)** T1w/T2w-FLAIR myelin maps from three representative equivolumetric layers (EVL). **(F)** PHT00 atlas (Paxinos et al., 2000) overlaid with area boundaries of the current study. **(D, E, I, J)** Sharp gradient-ridges vary across EVLs, suggesting areal boundaries. The area boundaries resemble those mapped by Hackett and colleagues. Therefore, we followed their areal/subareal nomenclature (Hackett et al., 1999, Kaas and Hackett, 2000). Abbreviations: AKL: auditory koniocortex, medial part; AKM: auditory koniocortex lateral part; Aud: primary auditory cortex; PAar: Paraauditory rostral; PaIM: Parainsular medial; ProK: Prokoniocortex; ProKL: prokoniocortex lateral; ProKM: prokoniocortex medial; ReI: retroinsular; REIp: retroinsular parietal; ReIT: retroinsular temporal (Paxinos et al., 2000).

Dorsal to the auditory cortex, the small densely myelinated “island” is a strong candidate for the parieto-insular vestibular area (PIV) (Pandya and Sanides, 1973, Grusser et al., 1990, Akbarian et al., 1994, Lopez et al., 2011, Guldin et al., 1998). The PIV exhibits high myelin content, which is likely important for swift transmission of neural signals related to 3D rotation and translation (Chen et al., 2010), key for maintaining postural stability. Vestibular-receiving areas are mostly distributed around the temporo-parietal junction, including areas 7a, 7b, 7op, SII and Tpt, as well as the posterior insula and parts of cingulate cortex (Guldin et al., 1998, Chen et al., 2010). Additionally, the subdivision of areas 3a and 3b in the upper limb region may correspond to the 3a–hand–vestibular area (3aNv; Supp. Fig. 4) (Guldin et al., 1998).

**Superior Temporal Cortex**

In the superior temporal cortex, we identified a total of 10 areas and numerous potential subdivisions (Supp. Fig. 6). In this region, myelin gradient-ridges were robust in several EVLs enabling relatively high-confidence determination of several area boundaries. Inspection of T1w/T2w-FLAIR maps and gradients in EVL3b reveals boundaries for the high and intermediate myelinated regions including MT/V5, MSTd, MSTv, V4t and TEOm (Supp. Fig. 6B,D). V4t was recognized by distinctive lighter myelination (Desimone et al., 1986). Between MT/V5 and MSTd a narrow homogeneous relatively lightly myelinated strip with sharp myelin gradients is identified as MTt (analogous to the name V4t: V4 transitory).

In the more sparsely myelinated areas, strong gradient-ridges in EVL5b reveal additional borders for areas STPc, DP, FST and an area with intermediate myelin content connecting the MT complex and auditory cortex. We use the term temporoparietal occipital junction (TPOJ) to describe the topological similarity of the area discovered in humans (Glasser et al., 2016). In humans, the intermediately myelinated region adjoining the MT complex and auditory cortex contains two functionally distinct areas TPOJ1 and TPOJ2 (Fig. 7B). It will be interesting to investigate whether the macaque TPOJ can also be functionally delineated into two separate areas and whether these parcels exhibit homologous functional connectivity with humans.

Comparison of myelin in EVL3b and EVL5b suggest further subdivision of area V4t whose inferior limits are uncertain: the inferior area/zone may correspond to the V4tp which has slightly sparser myelin than area V4t(a) (Lewis and Van Essen 2000).

Potential subdivisions were observed in the area MT/V5. Concentric layer-specific structures and a less myelinated ventral portion are notable in EVL3b (Supp. Fig. 6D). The MT is further divided in horizontal orientation revealed by midthickness-weighted myelin maps (Supp. Fig. 6I). The former subdivisions might correspond to areas MTp and the latter to MTc (Desimone and Ungerleider, 1986). The concentric layer-specific structures may be related to thalamocortical connections (e.g., outer band of Baillarger), whereas radial fibres could be related to intracortical communication pathways targeting specific superficial layers.

These distinct anatomical subdivisions in layer and column orientations might explain, in part, the numerous anatomical and physiological parcellations of area MT (Desimone 1986, Paxinos 2000, Lewis and Van Essen 2000). Numerous functions have been attributed to MT including direction selectivity (Maunsell and Van Essen 1983), speed sensitivity (Perrone et al., 2001), motion coherence (Britten et al., 1992, Rees et al., 2000), stereoscopic depth (DeAngelis et al., 1999), scotopic (low-light) vision (Ikeda et al., 2023), and eye movement control (Komatsu and Wurtz et al., 1988). The MT subdivisions may differentially contribute these functions, upper and lower visual fields, and receptive field properties, but there is not yet sufficient evidence to determine whether these gradient-ridges indicate distinct subdivisions, retinotopy, or distinct areas.

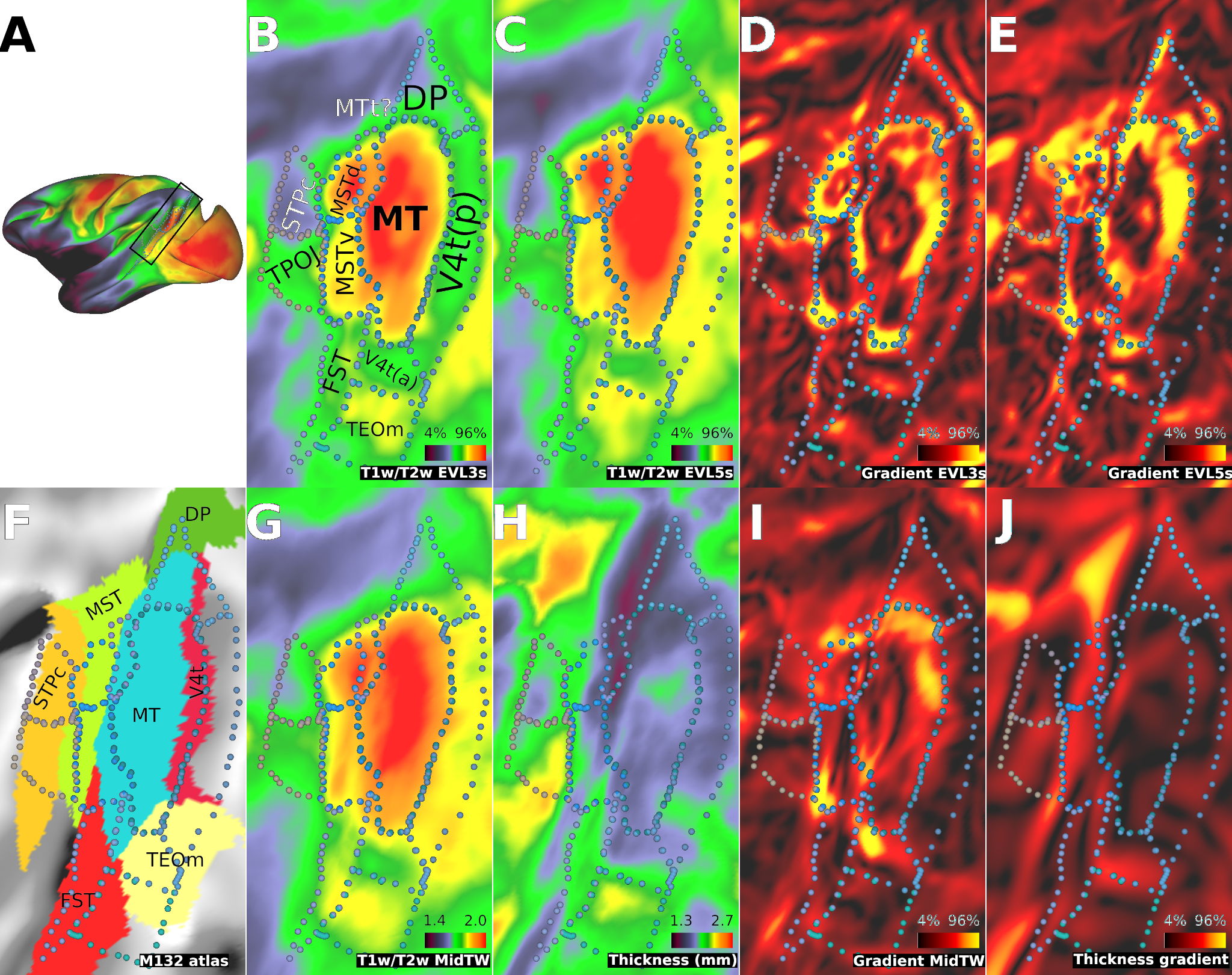

**Supplementary Figure 6. Charting Area Boundaries adjacent to MT complex (MT+).** **(A)** Location of the MT complex indicated by the snippet (black box). **(B, C)** T1w/T2w-FLAIR from two representative equivolumetric layers (EVLs) 3b and 5b. **(C, D)** Sharp gradient-ridges vary across EVLs, indicating areal boundaries. Two putative new cortical areas, or transition zones, are proposed: transitional MT (MTt) and temporo-parietal-occipital junction (TPOJ). The term “MTt” describes a narrow zone similar to V4t, while “TPOJ” refers to an intermediately myelinated area connecting the MT complex with the auditory cortex, previously identified in humans (Glasser et al., 2016). Abbreviations: DP: Dorsal prelunate (May et al., 1986); FST: Fundus of the superior temporal sulcus (Desimone and Ungerleider 1986); MT: Middle temporal area (Zeki 1974); MSTd: Dorsal medial superior temporal (Desimone and Ungerleider 1986); MSTv: Ventral medial superior temporal (Desimone and Ungerleider 1986); STPc: Caudal superior temporal parietal area (Bruce et al., 1981); TPOJ: Temporal parietal occipital junction (current study; in humans, see: Glasser et al., 2016); V4t: V4 transitional area (Desimone and Ungerleider 1986).

**Intraparietal Cortex**

In the intraparietal cortex, we identified a total of 14 areas/sub-zones (Supp. Fig. 7). In this region (Supp. Fig. 7A), myelin gradient-ridges were robust in several EVLs enabling relatively high-confidence determination of several cortical areas. Inspection of T1w/T2w-FLAIR maps and gradients in EVL3b and EVL5b reveals boundaries for the majority of the cortical areas (Supp. Fig. 7B-E). The areas LIPv and LOP were recognized by very dense myelin content and delineated using strong gradients (Supp. Fig. 7D). The areas AIP, MIP and LIPd were recognized by distinctively light myelin, however, the gradient between areas AIP and LIPd was weak and it was difficult to determine the precise location of the border between these two areas. Areas VIPl and VIPm (Lewis and Van Essen 2001) were recognized by sharp elongated gradients at the bottom of sulcus (Supp. Fig. 7I) with VIPl expressering lighter myelin relative to the surrounding areas (Table 1. in Lewis and Van Essen 2000). Area MDP was identified by its sparse myelin and V6A by its very dense myelin (Galletti et al., 1999, Lewis and Van Essen 2000). Area 5v exhibited heterogeneous myelin content (Supp. Fig. 7I), and subdivisions may exist in this area or the area VIPm extend to this region. Finally, zone 7t was identified as a small enclave at the anterior end of the intraparietal sulcus (data not shown).

Overall, we replicated several defining/distinctive factors and topological arrangement of cortical areas described in the LV00 macaque fascicularis monkey multi-modal histology parcellation (Supp. Fig. 7F) (Lewis and Van Essen 2000, Van Essen et al., 2012). Relative to the LV00 atlas, we recognized a putative new area which is a distinct intermediately/densely myelinated region within the area PO and between the areas V6A and MIP. The identity of this area is unclear: it may correspond to the V6 complex or to the area junction of parietal and superior temporal (JPS) discovered in humans (Glasser et al., 2016). Moreover, there is a thin strip of relatively high myelin between areas MDP and MIP, not described in previous atlases. Finally, we note that area LOP also topologically and by heavy myelination resembles the area V7 in humans (Tootell et al., 1998, Glasser et al., 2016), albeit advancing such hypotheses would require further functional investigations.

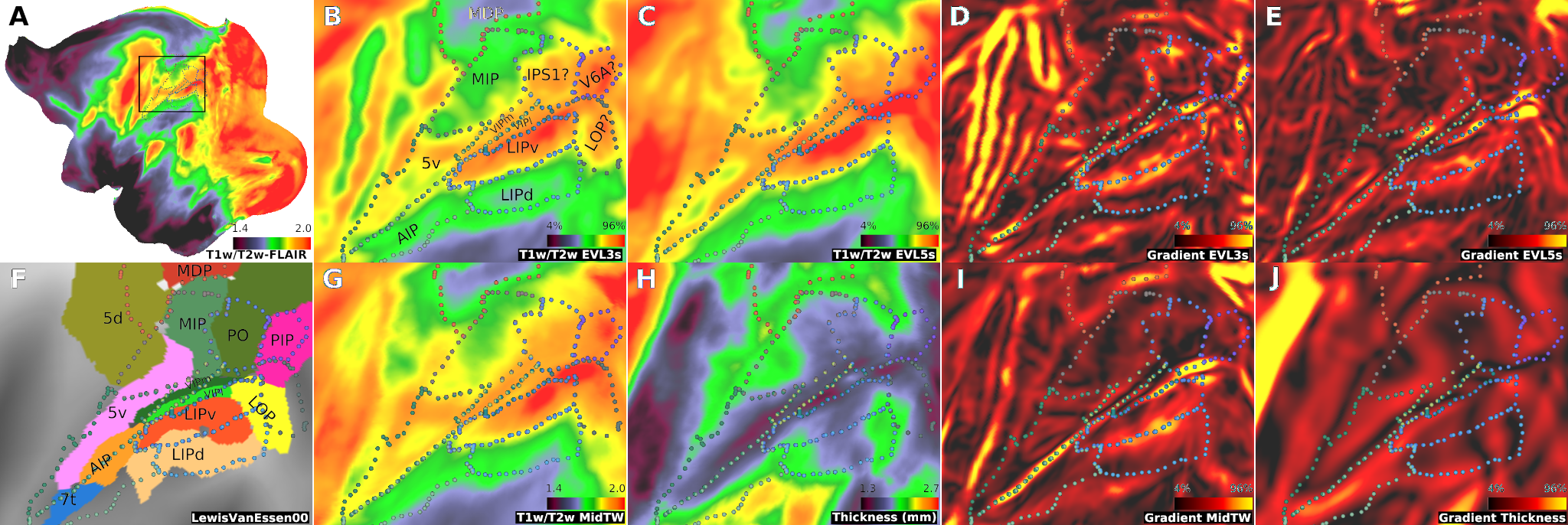

**Supplementary Figure 7. Charting Area Boundaries in Intraparietal Cortex.** **(A)** Location of the zoomed view indicated by the snippet (black box). **(B, C)** T1w/T2w-FLAIR from representative equivolumetric layers (EVL) 3a and 5b. **(D, E)** Sharp gradient-ridges vary across EVLs. **(F)** The area boundaries closely resemble those mapped by Lewis and van Essen (Lewis et al., 2001) with the exception of a potential discovery of an area JPS (Glasser et al., 2016). **(G, H)** Midthickness-weighted T1w/T2w-FLAIR, cortical thickness and **(I,J)** their gradients. Abbreviations: AIP: Anterior intraparietal area; (Preuss and Goldman-Rakic 1991); JPS: Junction of parietal and superior temporal (Glasser et al., 2016); LIPd: Lateral intraparietal dorsal area (Lewis and Van Essen 2000); LIPv: Lateral intraparietal ventral area (Lewis and Van Essen 2000); LOP: Lateral occipital parietal (Van Essen et al.,1983 ); MDP: Medial dorsal parietal area (Golby et al., 1988); MIP: Medial intraparietal area (Colby et al., 1988); PIP: Posterior intraparietal area (Cavada et al., 1989); PO: Parietal-occipital area (Andresen et al., 1990); VIPm: Ventral intraparietal medial area (Lewis and Van Essen 2000); VIPl: Ventral intraparietal lateral area (Lewis and Van Essen 2000); V6A: Visual area 6A (Galletti et al., 1999); 5v and 5d: Area 5 ventral and dorsal (Pandya et al., 1982).

**Orbitofrontal Cortex**

In the orbitofrontal cortex, we identified a total 14 areas (Supp. Fig. 8). This region was perhaps the most challenging region to parcellate due to the very weak myelin contrast (Supp. Fig. 8A). Layer-specific myelin gradients were negligible in EVLs1-4 and barely distinguishable in the deep EVLs 5a/b (Supp. Fig. 8B, D). On the other hand, midthickness-weighted myelin maps provided slightly stronger gradients, and the area boundaries were mainly determined using these gradients. (Supp. Fig. 8E). This result suggests that the small myelin differences between areas were relatively constant across cortical layers and the Gaussian-weighting procedure across the layers (see methods) improved the CNR between the cortical areas. Together, these results are compatible with neuroanatomy of agranular cortex (Fig. 6H): the agranular cortex is defined by the absence of layer IV, and weak/negligible bands of Baillarger (Carmichael and Price 1994). The vast majority of cortico-thalamo-cortical connections are in the cortical layer VI (Xiao et al., 2009), in contrast to sensory areas which contain the largest connections at layer IV. Taken together, the lack of high CNR layer-specific gradients may be attributed to the absence of bands of Baillarger.

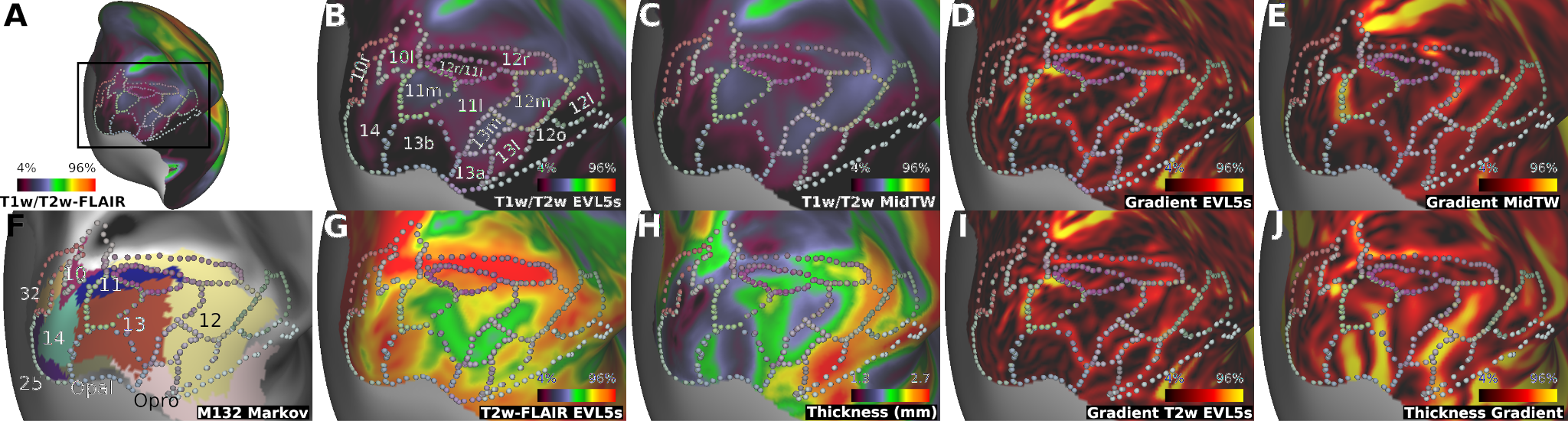

**Supplementary Figure 8. Charting Area Boundaries in Orbitofrontal Cortex.** **(A)** Anterior view of the cortex. The box indicates the location of the zoomed view. **(B)** Representative equivolumetric layers 5b (EVL5b) and **(C)** midthickness-weighted T1w/T2w-FLAIR. **(D)** In EVL5b, myelin gradient-ridges are relatively weak and **(H)** midthickness-weighted myelin gradients appear slightly more robust. **(F)** M132 atlas (Markov et al., 2011). **(G, I)** T2w-FLAIR and its gradients. **(H, J)** Cortical thickness and its gradients. Abbreviations Opal: Lateral opercular area; Opro: Opercular part of area Prol;

**Insular Cortex**

In the insular cortex, we delineated a total of 14 areas/sub-zones (Supp. Fig. 9). The dorsal “primary interoceptive areas”, Idfa and Idfp (Craig 1994, Craig 2002, Evrard 2014), were delineated by sharp myelin gradient-ridges in EVL5b in the dorsal fundus of the insular cortex (Supp. Fig. 9B, D). In the anterior granular insula, we delineated areas Igd and Igv (Ervard et al 2014). The layer-specific gradients between granular and dysgranular areas were strong in EVL5b which may be attributed to distinct distribution of thalamo-cortical fibres (Jones et al., 1976, Craig 1994).

In the dysgranular insular cortex (e.g., areas Idd, Idm, Idv, and Ivfp) myelin gradient-ridges became progressively weaker, and it was more challenging to demarcate subtle areal boundaries. We identified area Idd using strong-layer specific myelin gradients (Supp. Fig. 9D, E) and T1w/T2w-FLAIR peaks at EVL4b/EVL3b (Supp. Fig. 2A), consistent with Ervart and colleagues' description of a distinct outer band of Baillarger in sublayer IIIb (see their Fig. 7; Ervart et al., 2014). Area Idm was identified by relatively higher myelin in EVL6B (Supp. Fig 8C; Fig. 2C), relatively thick cortex (Supp. Fig. 9H) and absence of evident outer bands of Baillarger, which are also consistent with Evrart and colleagues’ description (see Fig. 8 in; Ervart et al., 2014). Note that this intermediately myelinated stripe very deep in the cortex is unlikely due to the partial volume of the white matter because this area has thicker cortex (2.6 mm) than neighbouring areas Idd and Idv (1.9 and 2.1 mm, respectively) and therefore has the smallest expected partial volume effect. Areas Idm, Idv and Ivfp exhibited myelin gradients in EVL6s and in midthickness-weighted maps (Supp. Fig. 9E, I).

Among agranular insular areas (e.g., Iapl, Iai, Ial, Iap, Iapm, and Iam), characterised by the absence of layer IV (Ervart et al., 2014), layer-specific myelin gradients were largely absent, and only border areas exhibited sharp layer-specific myelin gradients (e.g., Idm vs Iap; Supp. Fig. 9B, E). In contrast, midthickness-weighted myelin maps exhibited gradient-ridges, which were used to delineate areal borders (Supp. Fig. 9I). These weak areal differences appeared more pronounced in T2w-FLAIR images than in T1w and T1w/T2w-FLAIR images (data not shown), suggesting that the variations in MR signal-intensity may have originated from other factors than myelin, such as cell density, column/module density, or blood volume.

Overall, the topological arrangement of our 14-area partition in Macaca Mulatta resembles in several aspects the 15-area partition in the cynomolgus macaque monkey (Macaca Fascicularis) insular cortex by Ervart and colleagues (Ervart et al., 2014, 2019). The boundaries run along the posterior-anterior axis of the insular cortex and exhibit several layer-specific features previously described using histology. However, several areas exhibit inhomogeneity along the posterior-anterior axis, as Ervard and colleagues have previously noted. For instance, area Idd exhibits myelin inhomogeneity, and it is not clear based on myelin content alone whether these correspond to distinct areas or portions of an interoception/sensory homunculus. Further investigation of area borders using fMRI may help refine and elaborate the significance of these borders.

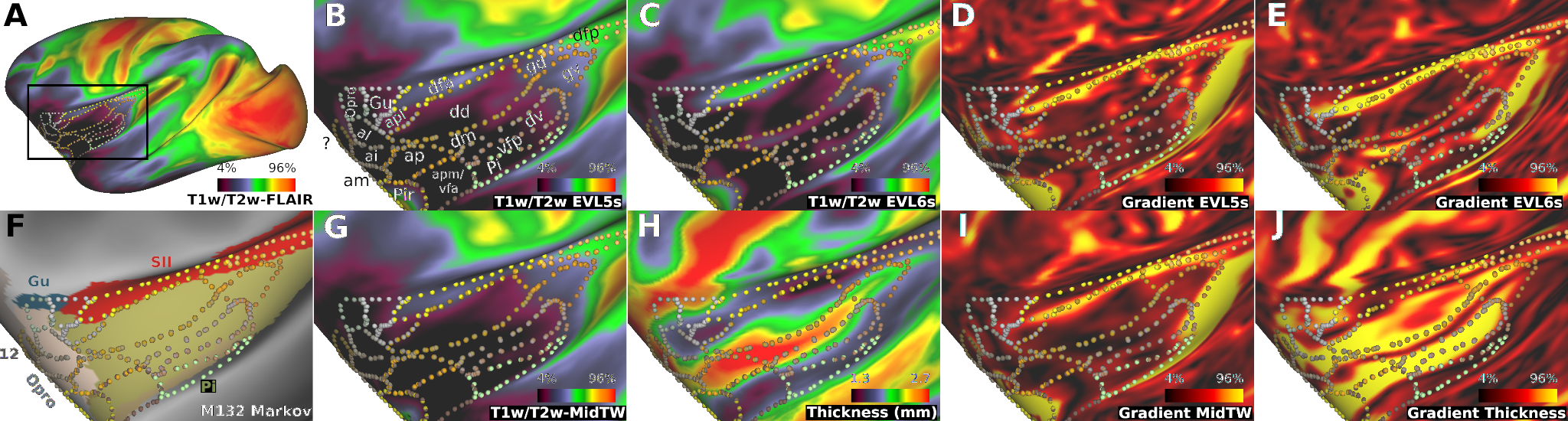

**Supplementary Figure 9. Charting Area Boundaries in Insular Cortex**. **(A)** Location of the zoomed view of the insular cortex is indicated by the black box. **(B, C)** T1w/T2w-FLAIR from representative equivolumetric layers (EVLs) 5b and 6b. **(D, E)** Myelin gradient-ridges are weak, due to sparse myelination, and only visible in the deep cortical layers. **(F)** M132 atlas (Markov et al., 2011) overlaid with area boundaries of the current study. **(G, I)** T2w-FLAIR exhibited putative area boundaries, potentially reflecting other architectonic factors such as cell density. **(H, J)** Cortical thickness and its gradients. Parcellation nomenclature followed those introduced by (Carmichael and Price, 1994) and (Ervard et al., 2013), albeit we could not distinguish all their areal boundaries and/or subdivisions (e.g., Iapm/Ivfa). Abbreviations: Gu: Gustatory cortex (Penfield and Faulk 1955); lai: Intermediate agranular insula (Carmichael and Price, 1994); Ial: Lateral agranular insula (Carmichael and Price, 1994); Iam: Medial agranular insula (Carmichael and Price, 1994); Iap: Posterior agranular insula (Ervard et al., 2013); Iapl: Posterior-lateral agranular insula (Carmichael and Price, 1994); Iapm: Posterior-medial agranular insula (Carmichael and Price, 1994); Idd: Dorsal dysgranular insula (Ervard et al., 2013); Idfa: anterior area of the dorsal fundus of the insula (Ervard et al., 2013); Idfp: posterior area of the dorsal fundus of the insula (Ervard et al., 2013); Idm: Mound dysgranular insula (Ervard et al., 2013); Idv: Ventral dysgranular insula (Ervard et al., 2013); Ivfa: Anterior area of the ventral fundus of the insula; Ivfp: posterior area of the ventral fundus of the insula (Ervard et al., 2013); Opro: Olfactory proisocortex; Pi: Parainsular area (Saleem et al., 2007); Pir: Piriform cortex (Saleem and Logothesis 2012).
